# Autonomous error detection is enabled by conflict-dependent forward models in human medial frontal cortex

**DOI:** 10.64898/2026.04.29.721757

**Authors:** Jake Gavenas, Zhongzheng Fu, Adam Mamelak, Ueli Rutishauser

## Abstract

Learning from mistakes is fundamental for survival. Humans can monitor their behavior to detect action errors without external feedback, utilizing error information for adaptation and learning. How such errors are detected remains unknown. We investigated single neurons recorded from human medial frontal cortex while subjects performed a task in which conflict between target and distractor increases control demands and error likelihood. Population activity in presupplementary motor area (preSMA), but not dorsal anterior cingulate, encoded target and distractor responses distinctly and only in conflict trials. These representations were insensitive to response accuracy, indicating they reflect forward predictive models. Geometric alignment between target and actual response representations covaried with error commission, supporting the mismatch theory of error monitoring. Artificial neural networks trained to detect errors developed similar neural geometries. These findings identify a mechanism for detecting errors without external feedback and show that all signals required for this computation are present in preSMA.

## Introduction

The ability to learn from mistakes is a cornerstone of human behavior. Remarkably, we can learn from mistakes even if we don’t receive external feedback informing us that we’ve erred. This is because humans can self-monitor and detect action errors as they occur. For instance, if you accidentally call your friend by the wrong name, you may realize your mistake as the incorrect name exits your mouth—you may even try to stop yourself from committing this faux pas or be more careful in future interactions. This process, often referred to as *internal or autonomous error monitoring,* is critical for rapid correction of behavior, adaptation, and learning ^1–6^. Over- and under-active error monitoring is thought to be a key aspect of OCD and schizophrenia, respectively ^7–9^, and disrupted error monitoring is at the core of an array of other psychiatric illnesses ^10–15^. Understanding the neural mechanisms of internal error monitoring is therefore important both for understanding how the brain controls behavior as well as for developing novel treatments for psychiatric illnesses. However, how the brain implements internal error monitoring remains unclear.

Error monitoring is thought to depend on the medial frontal cortex (MFC). This is because MFC encodes action errors ^14,16–19^ and individual MFC neurons signal errors as or immediately after they occur ^20–27^ (see also Fig 1E). Beyond signaling errors, MFC error-related activity is also causally related to post-error behavioral adjustments ^28^, indicating MFC’s central role in bridging error detection to adaptive behavior. Within MFC, action error signals emerge first in the presupplementary motor area (preSMA) followed shortly after in dorsal anterior cingulate (dACC) ^2,20^. These findings suggest that preSMA detects errors and transmits error information to dACC, which subsequently adjusts response parameters to improve behavior (for instance, by increasing response thresholds to improve accuracy, leading to post-error slowing) ^29^. While this body of work clearly establishes the existence of self-monitored error signals in preSMA and dACC (key parts of the MFC), what remains unknown is how these error signals are computed.

**Figure 1.**
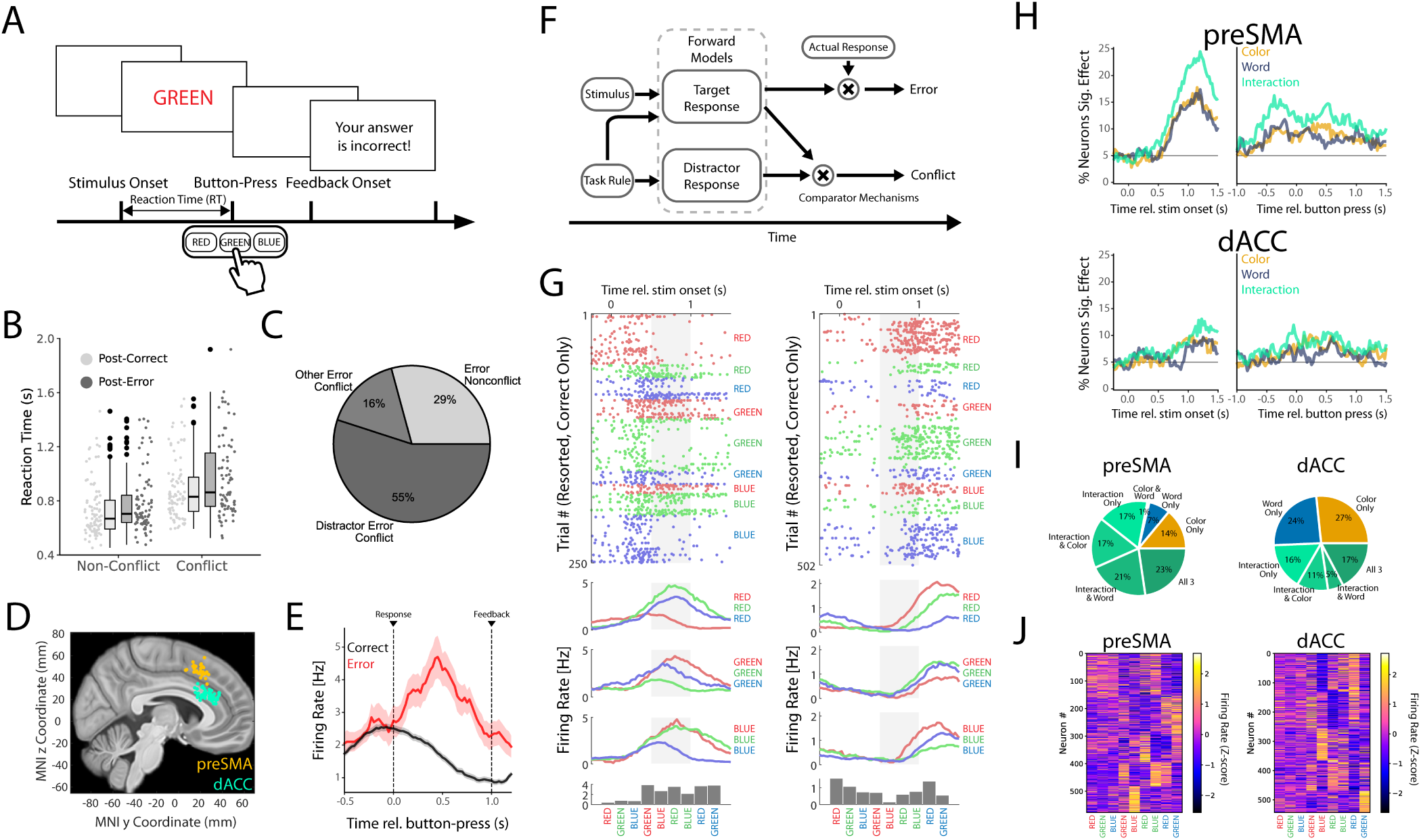
Task, behavior, and single-neuron results. (**A)** The task. Conflict emerges when the meaning of the displayed word and its ink color do not match. (**B-C)** Behavior. (**B**) Participants responded more slowly on conflict than non-conflict trials, with responses of both types slower if the preceding trial was an error (only correct trials shown). (**C**) Types of errors. Shown is the proportion of errors trials. In most error trials, subjects responded with the distractor, e.g. responding “Green” for the stimulus shown in (A). **(D)** Recording locations. **(E)** Example error neuron. Firing rates rapidly increase following error responses but not correct responses, prior to external feedback being given. (**F)** Hypothesized mismatch model (see text). (**G)** Two example preSMA neurons: raster plots (top), PSTHs (middle; 500 ms window; grey area is window used for statistical analysis of firing rates), and firing rate in the window [0.5 s, 1 s] following stimulus onset (“stimulus window”, bottom). **(H)** Proportion of neurons with significant tuning for color, word, and interactions (nonlinear mixing) between the two. Analysis is based on a 3×3 ANOVA with color and word as the main factors. **(I)** Proportion of neurons’ tuning profiles as a percentage of neurons that showed any kind of significant tuning in the window (main effect for color, word, or color-word interaction; in [0.5 s, 1 s], gray box in (G)) following stimulus onset. Nonlinear mixing was more common in preSMA than dACC. **(J)** Response profiles of all recorded neurons. Average firing rates for each of the 9 conditions (x axis) were calculated during the stimulus window and z-scored across conditions. Neurons with similar response profiles were grouped together using hierarchical clustering for visualization purposes.

The “mismatch” model of action error computation ^17,29–31^, an influential theory whose predictions remain largely untested, proposes that (i) forward predictive models ^32^ integrate task and stimulus information to predict the correct and incorrect response for a given situation, and (ii) action errors are detected by comparing forward model output (target response) with a copy of the motor command for the actual response (see Fig 1F). If this model explains how MFC detects action errors, representations of the predicted correct and actual response (the action) should be present in MFC. In contrast, alternative models make distinct testable predictions. They propose that errors are computed subcortically and then transmitted to MFC through dopaminergic pathways ^33,34^ or that error signals reflect amplified conflict signals or generic prediction error signals rather than the output of a target-response mismatch computation ^35–37^. Under these models, no signatures of forward models are predicted to be present within MFC.

In the present study, we investigated how the brain internally detects errors by testing the foundational predictions of the mismatch model of error monitoring. We examined activity of individual MFC neurons recorded while human subjects performed a 3-option Stroop task. We tested the hypotheses that (1) forward models and the actual motor response are separately represented in preSMA, and (2) that these representations are sufficient to detect errors through mismatch computations. In the Stroop task, it is well established that many neurons (Figure 1E shows an example) signal action errors rapidly after button-press, and well before feedback onset (Fu et al., 2019;). Peri-response error signals are also visible at the LFP or scalp EEG level as the error related negativity [ERN] ^2,17,18^, a widely studied non-invasive signature of error monitoring. This indicates that subjects can detect that they made an error internally in our task, without relying on external feedback.

## Results

### Task, Behavior, Single-Neuron Results

We investigated the activity of 1259 neurons recorded from two parts of the human medial frontal cortex: pre-supplementary motor area (preSMA, N=629) and dorsal anterior cingulate cortex (dACC, N=630). Recordings were obtained from patients (N=34, 87 sessions) undergoing epilepsy monitoring (see methods). Subjects performed a Stroop task with three colors (Fig. 1A): subjects were shown one of the words ‘red’, ‘green’, or ‘blue’ on a screen, displayed in either red, blue, or green font color. They were instructed to indicate the font color (referred to as target response or color) of the displayed word with a button-press, ignoring the meaning of the word (referred to as distractor response or word). Following a button press, the screen went blank and subjects were given feedback after a 1-second delay (Fig 1A). Subjects exhibited three hallmark behavioral findings of cognitive control expected in this task: (i) response times were longer when font color and word meaning disagreed, termed ‘conflict’, compared to when they agreed, termed ‘nonconflict’ (means ± standard error: 898±41 vs. 766±41 ms, p < 0.001; linear mixed model), (ii) response times in the trial following an error were slower than those following correct trials (850±40 ms vs. 810±4 ms, p < 0.001; linear mixed model; Fig 1B), and (iii) errors were more likely on conflict compared to nonconflict trials (7.9±0.8% vs. 5.5±0.5%, p < 0.001; logistic mixed effects; Fig 1C). What did subjects do when they made an error? The most common error in conflict trials was to press the button that corresponds to the meaning of the presented word (i.e. the distractor response, e.g. pressing “Blue” for when the word Blue is written in red ink; Fig 1C). This response pattern suggests errors were largely driven by failures of control rather than lack of engagement or attentional slips, in which case we would expect subjects to be equally likely to erroneously respond towards the third, uncued option as they were to the distractor.

Informed by mismatch models of error monitoring ^17,29,30^, we hypothesize that neurons in the MFC represent the output of forward models that compute the predicted target and distractor responses from stimulus and task information (Fig 1F), enabling the rapid detection of action errors (Fig 1E). A central prediction of this model is that the neurons that encode the target (font color) and distractor (word meaning) of a stimulus represent the outputs of the forward models needed to predict whether subjects made an error or not (Fig 1F) rather than representing stimulus information. To test this hypothesis, we examined the response of neurons in MFC (Fig 1D) separately for all possible word-color combinations.

Across all recorded neurons, response following stimulus onset (window [0.5 s, 1 s]) was significantly modulated by both color and word (example neurons in Fig 1G, S1). We examined single-neuron tuning for color and word using a 3×3 ANOVA with color and word as main factors. 15.3% and 9.2% of neurons in pre-SMA and dACC were significantly modulated by color, respectively (Fig. 1H; p<0.05 for main effect of color; proportions significantly larger than chance p < 0.001, binomial test). Similarly, 14.4% and 7.9% were significantly modulated by word identity, respectively (Fig. 1H; proportion significantly larger than chance p < 0.001 and p = 0.006, respectively).

Firing rate modulations were heterogenous across neurons, partially because interactions were prominent: the response of a significant portion of neurons was modulated by nonlinear interactions between word and color (21.9% in preSMA, p < 0.001; 8.3% in dACC, p = 0.002; Fig 1H). Notably, such “mixed selectivity” was especially prominent in preSMA, in which 78% of neurons that had either a main effect and/or an interaction exhibited nonlinear mixing (Fig 1I), an effect also apparent in the example neurons shown (Fig 1G) and the population (Fig 1J). These interactions are only partially explained by conflict (i.e. disagreement between color and word), which is a specific type of color-word interaction: in a combined ANOVA (3×3 color-word, with conflict as a specific covariate), 23.6% of preSMA cells exhibited either conflict or color-word interactions. Of these, 45.2% were ‘pure conflict’ (conflict with no significant other color-word interaction; Fig 1G left), 28.6% were ‘pure interaction’ (no effect of conflict, but a significant other color-word interaction Fig 1G right), and 26.2% exhibited both types of signals (see also Fig 3). This data shows that neurons in MFC encoded the hypothesized outputs of the two forward models, particularly in the preSMA. The output of the two forward models was, however, strongly intermixed, resulting in a highly heterogenous representation at the single-neuron level.

### Cognitive forward models in presupplementary motor area

Nonlinear mixing at the single-neuron level of the kind we found in our data (Fig. 1H) can, in some instances, result in linearly separable representations at the population level ^38,39^. We next tested whether this was the case for the representations of color and word in MFC. To do so, we formed *pseudopopulations* of all recorded neurons by aggregating time- and condition-matched neuronal responses (see Methods). We then trained decoders to differentiate between pairs of stimuli that differed in only one color or word feature (e.g. decoding BLUE vs BLUE to decode color; Fig 2A). We initially used only conflict trials for this analysis to avoid confounding our analysis with conflict signals (but see Fig. 3). Both the mismatch ^29^ and conflict-amplification ^40,41^ models predict that color (target response) is encoded, but only the former predicts that word (distractor) is also encoded.

**Figure 2.**
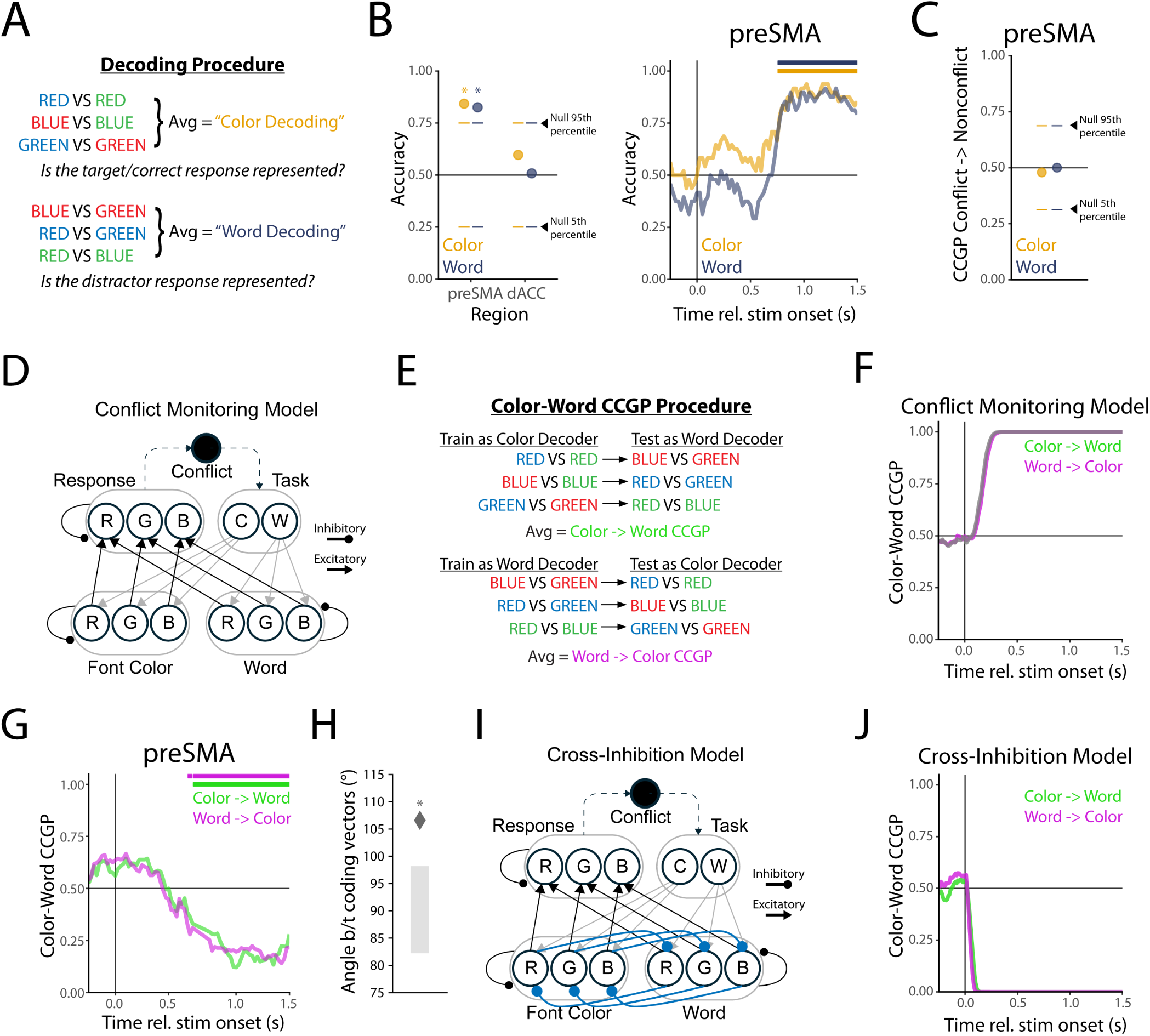
Neural geometry of color and word representations. (**A)** Decoding approach. We trained decoders to differentiate between condition pairs that differed along only one color or word. (**B)** Color and word information is decodeable from preSMA but not dACC. Horizontal bars mark periods of significant above-chance decoding (above 95^th^ percentile (one-sided test) of a null distribution obtained from bootstrapped decoders trained on data with shuffled labels, 1000 bootstraps). (**C)** At-chance CCGP of Color and word decoders trained on conflict trials to non-conflict trials. Null distribution calculated as in panel B. (**D)** The Conflict Monitoring Model. Color and word information activate corresponding nodes in the input layers. Those nodes in turn activate corresponding nodes in the response layer. Task representations bias the competition towards a desired result, and competition between response options gives rise to conflict signals (top node). (**E)** Color-Word CCGP procedure. We assessed whether color decoders generalized to differentiate between pairs of words mentioning the same colors and vice-versa. (**F)** Color and word decoders trained on simulated data from the conflict monitoring model exhibited high CCGP. (**G)**Color and word decoders trained on empirical preSMA data exhibited significantly *below-chance* CCGP (anti-generalization; left). Significance was assessed by comparing decoding results to the 2.5^th^ percentile of a null distribution (two-sided test) constructed as in panel B. **(H)** Color-word anti-generalization emerged because analogous color and word coding vectors had angles significantly greater than 90°. **(I)** Cross inhibition model schematic. We hypothesized that introducing lateral inhibition connections between analogous color and word nodes in the conflict monitoring model could serve as a mechanism for our results. **(J)** Color and word decoders trained on simulated data from the lateral inhibition model exhibited below-chance CCGP.

We found that both color and word were decodeable from neural activity in preSMA in single trials with an accuracy of 84.3% and 82.5%, respectively (Fig. 2B; chance is 50%; support vector binary classifier with linear kernel, stratified leave-two-out cross validation accuracy; spikes counted in window [0.5 s, 1.0 s] rel. to stimulus onset, p < 0.05 based on 1000 bootstrapped decoders trained on data with shuffled labels). In contrast, neither color nor word were significantly decodable from neural activity in dACC, with accuracies of 59.7% and 50.8% respectively (Fig. 2B, p > 0.05), was significantly higher in preSMA compared to dACC (Fig 2B, p < 0.001, rank-sum test; Cohen’s D = 1.89, 2.06 for color and word decoding respectively). Decodability of both color and word emerged at similar points of time in preSMA (Fig 2B, right, all color & word splits in Fig S2).

Color and word decoders relied on the neurons with single-neuron tuning in preSMA, as shown by significant correlations between decoder loadings and single-neuron F statistics (Fig S3). Remarkably, the neurons that decoders relied on the most were those with non-linear interactions (r=0.49 & r=0.52 for color and word decoders, both p < 0.001), with correlations weaker for the main effects (r=0.34 color, r=0.45 word, both p < 0.001). Suppporting this conclusion, removing neurons with strong interactions from the pseudopopulation rapidly reduced color and word decoding accuracy, whereas removing neurons randomly left decoding intact even for high numbers of removed neurons (Fig S4). This analysis shows that the non-linear mixed representations at the single-neuron level gave rise to linearly separable representations of color and word in the preSMA. In contrast, although there were single neurons in dACC that coded for color and word (Fig. 1H), at the population level color and word were not significantly encoded (Fig. 2B). For this reason, the analysis that follows focuses on the preSMA unless otherwise specified.

Why was color and word decodable from preSMA? We hypothesized that forward models compute the predicted target and distractor. However, a straightforward alternative interpretation is that our decoding results reflect stimulus representations rather than forward models. If this were the case, decoders trained on conflict trials should generalize to non-conflict trials that share the same stimulus features (cross-condition generalization performance, or CCGP; e.g. train a color decoder on BLUE vs BLUE, and test it on RED vs GREEN). If the identified representations are sensory, CCGP should be above chance due to the same stimuli features being present on conflict and non-conflict trials. We found that decoders did not generalize significantly to non-conflict trials, with average CCGP of 47.9% and 50.0% for color->word and word-> color respectively (p>0.05 based on 1000 bootstrapped decoders; Fig 2C). This result suggests that the representations we identified are not stimulus-based but rather the result of forward models which operate differently on conflict and non-conflict trials.

We next investigated the geometry of color and word representations in the preSMA. We used the Conflict Monitoring Model of the Stroop task as a guiding framework for generating predictions for the expected geometry ^35,36,42^ (schematic in Fig 2D). In the conflict monitoring model, stimulus color and word information activate congruent nodes in a response layer with winner-take-all dynamics, with task information biasing the competition so that the desired output is selected (Fig. 2D). Within this framework, a possible interpretation of our color and word decoding results is that preSMA encodes the response layer, with color decoding reflecting the feature being amplified (Fig 2D).

In geometric terms, by virtue of color and word information reflecting the same kind of underlying representation in the response layer, this model thus predicts that color and word information is organized along the same direction in neural state-space (that is, the coding directions are parallel to each other). If this is the case, then decoders trained to differentiate between one pair of colors (e.g. BLUE vs BLUE) should exhibit high CCGP when tested as a word decoder on the matching pair of words (RED vs GREEN). In contrast, at-chance CCGP would suggest a lack of a systematic relationship between the representations of color and word (which would be at odds with the conflict monitoring model). We confirmed this prediction in a simulation of the conflict monitoring model, in which, as expected, color and word decoders exhibited high CCGP (Fig. 2F; simulated data in Fig. S5A). This high CCGP in the simulated data was driven by the response layer: when restricting decoders to the input color and word layers, CCGP was at chance (Fig. S6A), confirming that color and word representations outside of the response layer are unrelated.

We next repeated the same analysis for the preSMA data. This revealed that decoders trained on pairs of colors and tested on corresponding pairs of words, and vice-versa, exhibited CCGP levels significantly below chance (Fig. 2G): Color->Word CCGP was 15.9% and Word->Color CCGP was 17.5% (both significantly below chance of 50% with p < 0.05 based on 1000 bootstrapped decoders trained on data with shuffled labels). Below-chance CCGP indicates that decoders “anti-generalize”, meaning that the binary decoders predict the opposite of the correct label rather than being at chance. For example, a red vs. green color decoder trained to differentiate between BLUE vs BLUE, when applied as a word decoder, consistently outputs the label “red” for GREEN trials and “green” for RED trials. This finding suggests that color and word representations are systematically related to each other in that they have parallel coding vectors (and coinciding separating hyperplanes), but with the coding vectors for related pairs pointing in *opposite* directions (Fig 2G). Compatible with this interpretation, the angles between corresponding color and word decoders (their coding vectors) were significantly greater than 90° (average angle = 106.6°, p < 0.05 based on shuffled data; Fig 2H; Fig. S7 for individual pairs). This organization is visible in low-dimensional representations of preSMA activity (Fig S8). These results indicate that unlike predicted by the conflict monitoring model, color and word encoding in preSMA do not reflect either unrelated representations or the same underlying representation.

What neuronal mechanism could give rise to the representational geometry we observed in preSMA? We reasoned that this geometry could be the result of competition between pairs of neurons representing the same color and word meaning, mediated through targeted inhibitory connections between those neurons (Fig 2H, bottom). This “cross-inhibition” model is motivated by our finding that representations of color and word are distinct on conflict trials (Fig 2C), in which a stimulus can never have the same color and word meaning. We developed a modified version of the conflict monitoring model with cross inhibition between corresponding color and word nodes to test this prediction (Fig 2H). These connections lead to suppressed activity in the color and word nodes corresponding to the cued word and color (e.g. presenting “blue” written in red ink results in relatively suppressed activity in the blue color and red word nodes; see Fig S5B). In simulated data from this version of the model, we observe significantly below-chance CCGP akin to the empirical data (Fig 2I). As hypothesized, anti-generalization was driven by the input layers in the cross-inhibition model: when restricting decoders to the response layer in the simulated data, above-chance CCGP remained (Fig S6B). Taken together, this model presents a plausible circuit-level mechanism for how forward models could give rise to the neural geometry we observed in preSMA.

### Conflict signals regulate forward model activity

Representations of conflict are a prominent feature of MFC, including at the single-neuron level ^21,36,43,44^, and theories of conflict detection suggest conflict signals are derived from color and word representations ^35,36,42^. We therefore next investigated whether the forward models we identified are related to conflict.

The responses of 23.7% of preSMA neurons (Fig 3A; p < 0.001, binomial test) were modulated by conflict in the stimulus period ([0.5 s, 1 s] relative to stimulus onset; Fig 3A), and conflict was highly decodeable from preSMA population activity, with an accuracy of 95.5% (firing rates in the window [0.5 s, 1 s] relative to stimulus onset; Fig 3B). Conflict signals in preSMA exhibited high CCGP between different pairs of stimuli (Fig 3C): conflict decoders trained leaving out one condition pair (e.g. RED vs BLUE) generalized to those held-out pairs (mean±SEM CCGP 84.7±0.28%, p < 0.05 based on bootstrapped distribution), indicating the overall dimension of conflict was invariant across specific stimuli. Accordingly, conflict coding vectors defined by individual condition pairs were highly similar with an average parallelism score of 0.356 (SEM=0.0007, p < 0.05 based on bootstrapped distribution; Fig S9). These results show that preSMA represents conflict as an abstract variable ^45^.

**Figure 3.**
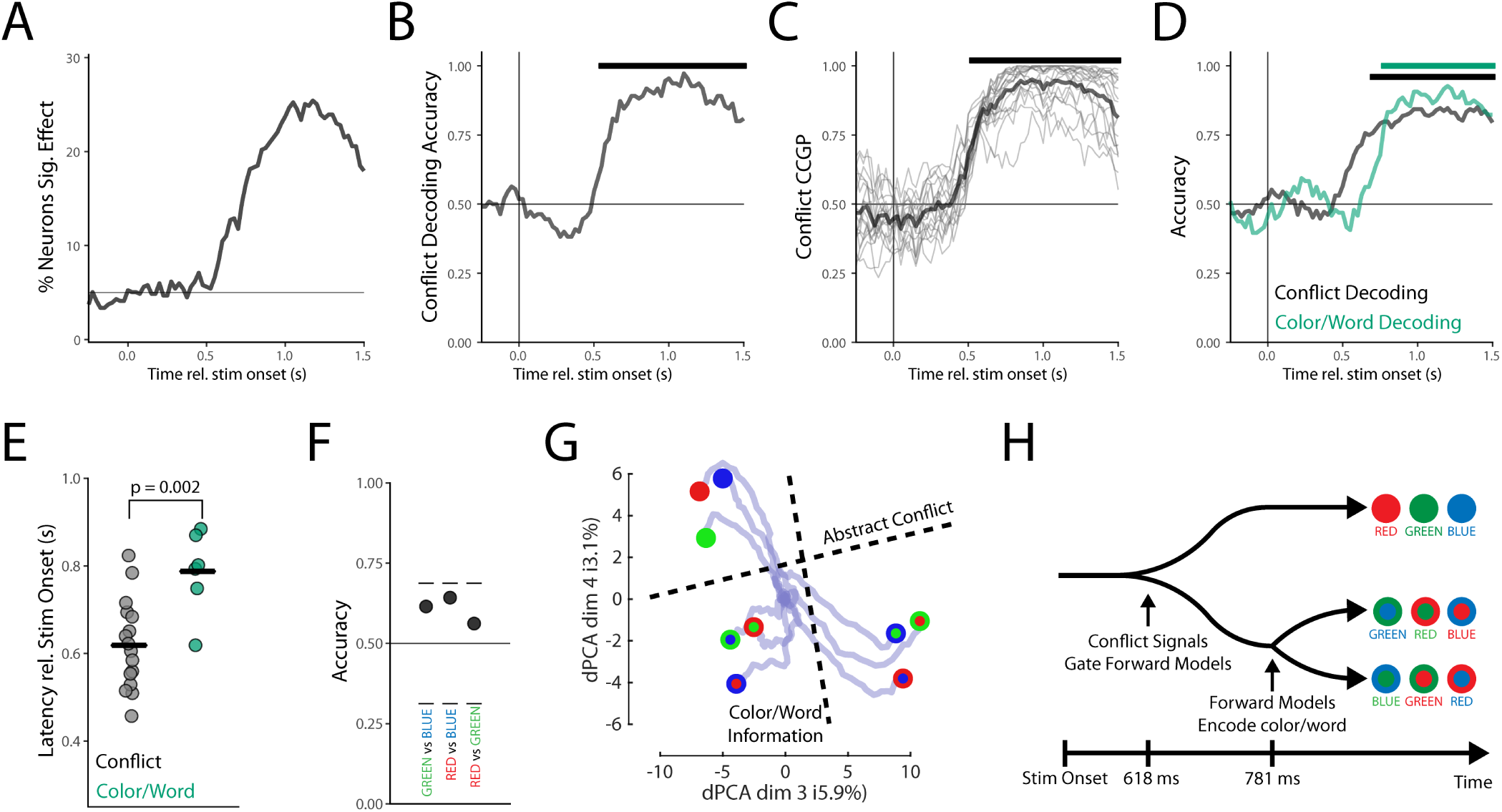
Hierarchical representational structure of conflict and forward models. **(A)** Proportion of neurons whose firing rates are significantly modulated by conflict (ANOVA). **(B)** Cross-validated conflict decoding accuracy from preSMA (black trace is average over cross-validations; black bar indicates significance, greater than the 95^th^ percentile of null distribution). **(C)** Conflict is abstract, with high CCGP of decoders trained on single pairs and trained on all others. Black line is average, grey lines are individual pairwise conflict decoders’ CCGP to all other pairs. Horizontal bars note period of significant above-chance CCGP. **(D)** Performance of different groups of pairwise decoders. Decoded pairs could be either one conflict and one nonconflict condition (conflict signals), two conflict conditions (color/word decoding as well as other possibilities), or two nonconflict conditions. **(E)** Conflict decoding became significant earlier in time than color/word decoding. Latency is relative to stimulus onset. **(F)** Nonconflict trials are not significantly decodeable from preSMA activity, implying conflict gates forward model activity. Points are average decoding across cross-validations, horizontal lines demark the 5^th^ and 95^th^ percentile of the null distribution. (**G)** Representational structure over time visualized with dPCA. Interior color reflects stimulus font color, and exterior color reflects stimulus word meaning. Point locations are at 1 second post-stimulus, and the blue lines reflect their trajectory since stimulus onset, where they began at the origin. The primacy of conflict signaling is reflected in the vertical orientation of the trajectories, showing that conflict emerges first in neural state space. **(H)** Schematic of hierarchical representations emerging across time. Conflict signals emerge first, and prompt preSMA to engage forward models, leading to the color-word geometry we identified previously.

The conflict monitoring model predicts that conflict is computed from color and word representations (Fig. 2D), which necessarily entails that conflict signals emerge after color and word signals. In contrast to this prediction, in preSMA conflict was decodable earlier than color and word meaning (Fig 3D). We quantified this effect by identifying the first point of time at which information was significantly decodable for individual conflict decoders (trained on specific pairs of conflict and nonconflict trials) and forward model decoders (trained as described in Fig 2). This analysis confirmed that conflict signals emerged significantly earlier than the forward models in preSMA (Fig 3E; p = 0.0024, two-sided independent samples t-test).

If conflict is not computed from color and word meaning, then how do conflict signals relate to error processing? We hypothesized that conflict may regulate forward model activity due to the increased likelihood of errors on conflict trials. This hypothesis echoes findings that MFC conflict signals regulate subsequent processing ^36,46–49^. To test this hypothesis, we investigated whether color and word were decodeable in non-conflict trials, because in non-conflict trials, color and word meaning are identical to each other, this reduces to asking whether pairs of words printed in their corresponding color are decodable from each other. Surprisingly, non-conflict conditions (e.g. RED vs GREEN) were not significantly decodeable from each other despite greater trial numbers than conflict trials, reaching an average binary decoding accuracy of only 60.6% (597 preSMA neurons, 16 trials in each condition compared to 8 in each condition for conflict trial decoding; using activity in the window [0.5 1] s rel stim onset; p > 0.05 compared to 95^th^ percentile of null distribution; Fig 3F). This analysis shows that the representations associated with color and word meaning are not present in non-conflict trials. This finding suggests that preSMA forward models are regulated by the detection of conflict, becoming active only after conflict has been detected in the first place.

To further characterize the relationship between conflict detection and forward models, we investigated neural activity using dPCA. We decomposed time-varying activity into task-defined components corresponding to color, word, and color-word interactions (some of which reflect conflict). Interaction components explained 29% of the neural variance as opposed to 11% and 10% for color and word respectively. The largest two interaction components correspond to conflict and the color-word representational structure we observed previously (Fig 3G, Supplementary Video 1). These findings indicate a dynamically changing representational geometry: abstract conflict signals emerge first, followed by further structure in which the six conflict conditions are separated into two groups such that stimuli of opposing color/word combinations are in separate groups (Fig 3G; schematic in Fig 3H). In our data, this second step is what led to the anti-generalization of color and word decoders. Strikingly, these phenomena can be observed at the single-neuron level: the two example neurons in Figure 1G show a neuron (left) that exhibits similarly high firing rates for all conflict trials, and lower activity for non-conflict trials. This neuron responds earlier than a color-word interaction neuron (right) that demonstrates a color-word inversion effect: firing rates are higher for RED than RED trials, yet higher for BLUE than GREEN trials. Taken together, these results indicate that preSMA forward models are regulated by conflict signals with conflict signals preceding the forward models.

### Forward models support error monitoring and detection

We next asked whether the forward models identified in preSMA (Fig 4A) inform cognitive control and/or performance monitoring. As a first step, we examined whether the same neural geometry is present in error trials (the analysis so far only included correct trials). We reasoned that if the forward models are utilized for controlling response selection in each trial, the neural geometry should be absent or distorted in cases of control breakdowns, namely on error trials. Conversely, if utilized for performance monitoring but not control, the representational structure must be preserved in error trials. To examine this question, we trained color and word decoders on correct trials from sessions with at least 6 errors in conflict trials and tested how well decoding worked in error trials (see Methods). In correct trials, color and word information was decodeable with high accuracy (Fig 4B, left; 93.8% and 91.7%, respectively). When tested on error trials, decoders generalized well with CCGP of 76.3% and 79.1% for color and word decoding, respectively (Fig 4B, middle; both p < 0.05, nonparametric bootstrapping). Furthermore, decoders trained on correct trials exhibited below-chance CCGP (anti-generalization) when generalizing from color to word and vice-versa on error trials (Fig 4B, right; CCGP 15.7% and 25.9% respectively, p < 0.05 both). This analysis shows that the forward models were also present in error trials, with a representational structure identical to that in correct trials. This indicates that these representations could subserve performance monitoring, a prediction we tested next.

**Figure 4.**
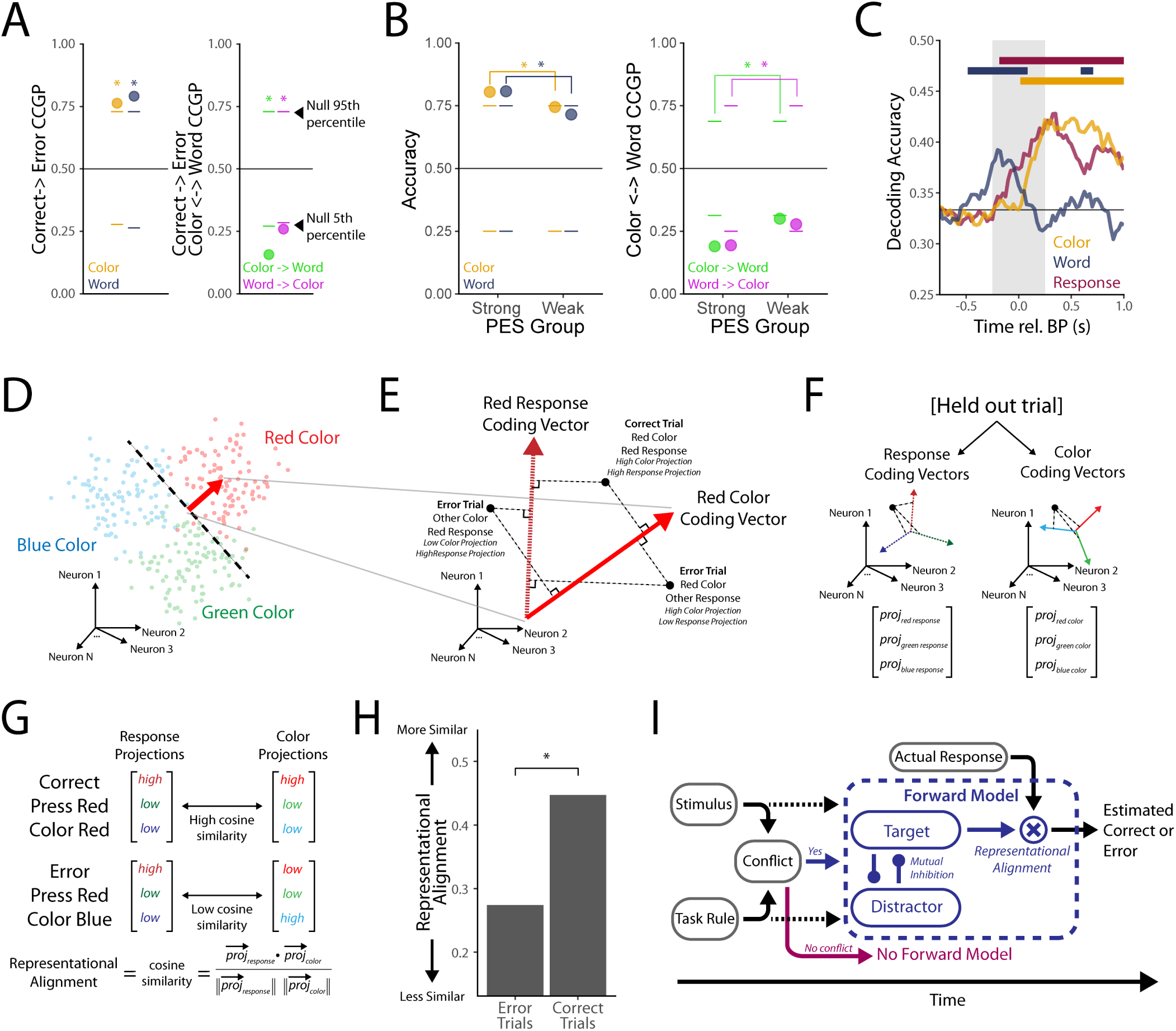
Role of forward models in error monitoring. **(A)** Decoders trained on correct trials from sessions with at least 6 error trials generalize to error trials (middle) and exhibit anti-generalization when generalizing from color to word and vice-versa (right). **(B)** Color and word decoding is higher, and anti-generalization is stronger in sessions with greater post-error slowing. **(C)** Color, word, and response can all be decoded around response onset. Horizontal bars reflect periods of significantly above-chance decoding according (assessed using independent samples t-test of decoding distribution against null distribution, constructed by repeating the procedure with shuffled labels; p < 0.05 FDR corrected for multiple comparisons across time). **(D-G)** Representational Alignment schematic. **(D)** Specific color and response (not shown) coding vectors were defined using a one-versus-rest procedure. **(E)** Considering matching pairs of color and response coding vectors, correct trials for that color/response pair will have high projections onto both coding vectors, whereas error trials will have low projections onto at least one of the coding vectors. **(F)** We used a cross-validated approach, projecting activity from held-out correct or error trials onto coding vectors defined using all other trials. **(G)** We define the representational alignment as the cosine similarity of the projections onto response and color coding vectors. For correct trials, alignment should be high due to a match of color and response, whereas for error trials, alignment should be low due to mismatched color and response. **(H)** Representational alignment is significantly higher for correct than error trials, indicating this measure could serve as the basis for error detection. **(I)** Updated schematic model. Conflict signals are computed *first* and then utilized to generate a combined target and distractor signal. This signal is then utilized alongside the actual response representation to compute errors via representational alignment.

A signature of successful performance monitoring is post-error slowing (PES), which is a reduction in response speed in trials that follow errors. We therefore next tested whether the forward models were related to PES by grouping sessions into those with strong and weak PES (median split; see methods). Comparing the neural representation between the two groups revealed that both the decoding and CCGP signatures of forward models were stronger in strong compared to weak PES sessions (Fig 4C; S10): strong PES sessions had greater color and word decoding at accuracy of 80.5% and 80.8% respectively (p < 0.05 both), compared to weak PES sessions’ respective accuracies of 74.5% and 68.8% (both p > 0.05; differences p < 0.001 both, rank-sum test; Cohen’s D = 0.43, 0.71 for color and word decoding respectively). We found similar results for color-word CCGP: anti-generalization was stronger in strong PES sessions (19.0% color->word and 19.4% word->color, p < 0.05 both) compared to weak PES sessions (30.0% color->word and 27.8% word->color, both p < 0.05; differences p < 0.001 both, rank-sum test; Cohen’s D = −1.03, −0.82 for color->word and word->color CCGP respectively). This data supports the view that the forward models subserved performance monitoring in a behaviorally relevant manner.

### preSMA detects errors via alignment of color and response representations

The mismatch model holds that the color (target) and/or word (distractor) representations provided by the forward models are compared with the actual response to predict whether subjects made an error or not without relying on external feedback (see Fig 1F). We next tested this model prediction by decoding both the target and actual response (the button pressed by the subject) and comparing those representations to investigate whether a preSMA could implement a mismatch computation. To decorrelate target and actual response (which are equivalent on correct trials, by definition) we performed this decoding analysis on a combined pseudopopulation constructed from both correct and error trials (see methods for pseudopopulation construction and decoding details).

Color, word, and response information were all decodeable from preSMA activity around response onset (Fig. 4C, S11), with word (distractor) information peaking first at an accuracy of 39.2% 0.2 s before button-press (chance level is 33%), followed by the emergence of color (target) and response button identity which reached peak decoding accuracy of 42.3% 0.525 s after button-press and 42.8% 0.350 s after button-press, respectively (chance is 33%; p < 0.001 all; see Methods). Response was also decodable in dACC, indicating this information may emerge due to a broadly-distributed signal (Fig S12).This data shows that in addition to the output of the forward models, response identity (also referred to a corollary discharge or an efference copy by others) is represented in preSMA, as has been predicted but never shown before ^17,22,29,50^.

These results show that preSMA encodes, on single trials, all the information required to generate an error signal in the way the mismatch model predicts. What remains to be determined is how the actual mismatch operation could be implemented. Some prior models of how the mismatch computation could be implemented rely on distinct neurons coding for color and response, for instance through AND and XOR gates ^29^. In our data, however, color and word information were best separable at the population level, with prominent mixed selectivity at the single neuron level. We therefore sought to determine how the mismatch computation could be implemented at the population level, culminating in an account we term Representational Alignment. We first define coding vectors for specific colors and responses using a one-versus-rest approach (binary decoders, such as ‘red vs. blue/green’, see Fig 5D). There are three such coding vectors for color, and three for response, each of which thereby define a 3-dimensional subspace (Fig. 5F). We then projected activity from held-out trials on all response and color-coding vectors, effectively into a response subspace and color subspace. In correct trials, color and response are equivalent, resulting in high projection values for matching pairs of color and response coding vectors (e.g. red-color and red-response, see Fig. 5E). In contrast, in error trials, projection magnitudes will be low for either the color or response coding vector for a given pair (Fig 5E). Therefore, comparing the three projection values between the color and response subspace implements the mismatch operation (Fig 5F). We operationalized this idea by defining representational alignment as the cosine similarity between the two projection value vectors (Fig 5G), with high and low cosine similarity indicative of correct and error trials, respectively.

**Figure 5:**
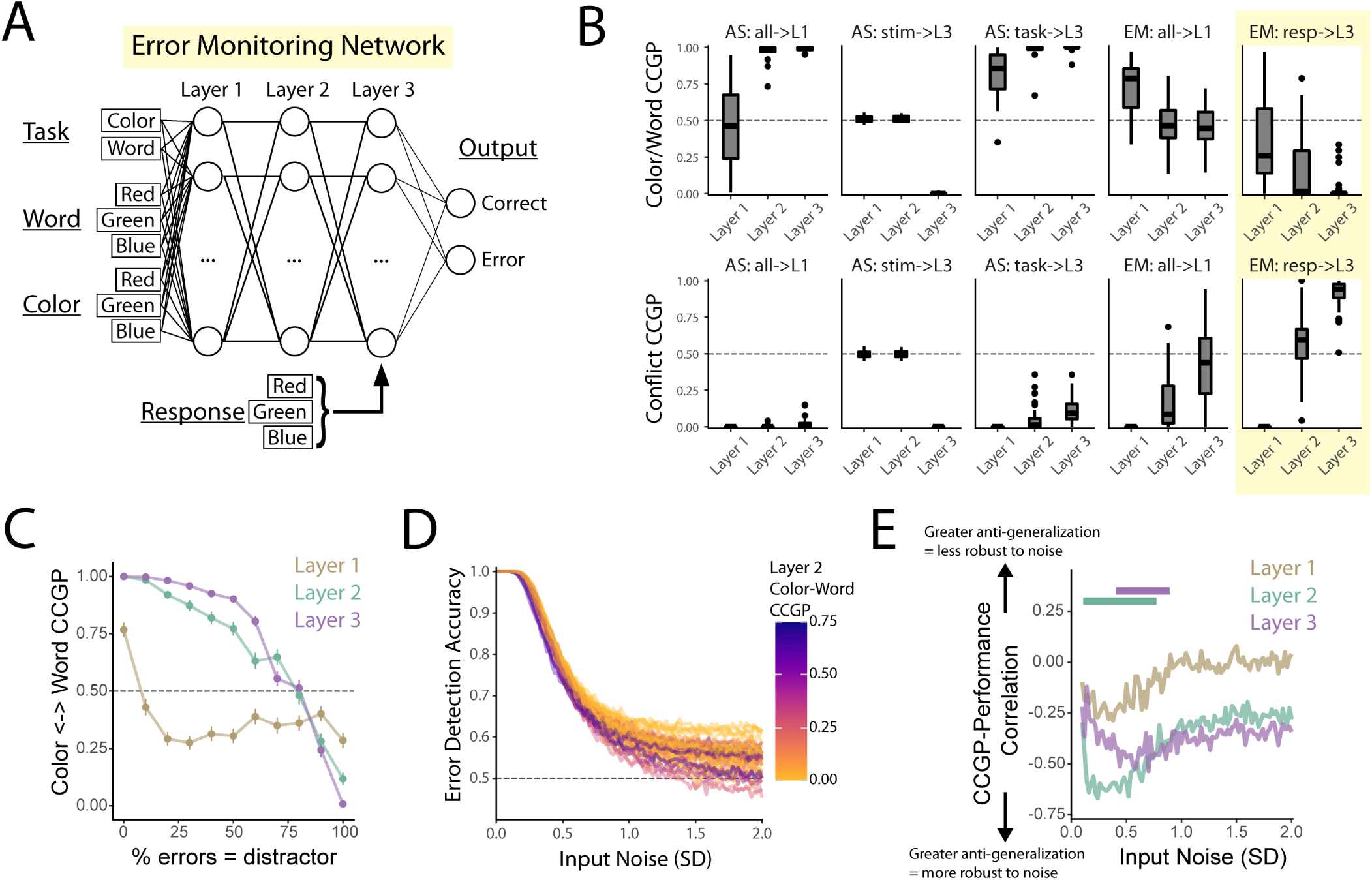
Feedforward neural network modeling of control and error monitoring in the Stroop task. **(A)** Stroop error monitoring network architecture. Task, word, and color are given to layer 1, and response information is given to layer 3 (all one-hot encoded). All layers (50 neurons each) are fully connected and use ReLU activation, with the final layer utilizing an additional softmax activation. (**B)** We trained 50 distinct networks for each of 5 distinct architectures and then conducted the color/word decoding and CCGP approach for each of the 3 layers separately (see Fig 2). Only the error monitoring network described in panel A (highlighted) consistently exhibited both color-word anti-generalization (top) and high conflict CCGP (bottom). (**C)** We tested whether the statistical relationship between the distractor response and error likelihood affected networks’ representational geometry. We parametrically varied how frequently error responses in the network matched the distractor versus the third, uncued, response option. Shown is color-word CCGP at all three network layers (means & standard error across 50 trained networks). Color-word anti-generalization was present only when most errors matched the distractor response, especially for layers 2 and 3. (**D)** For the error monitoring networks described in panel A we simulated activity with increasing noise, and found that networks were at first somewhat robust, and then performance sharply dropped off with increased noise. Each line reflects a network’s performance at different noise levels, colored according to their color-word CCGP. Networks that had stronger anti-generalization at layer 2 (before response signal delivery) seemed to exhibit more robust performance. (**E)** We correlated color-word CCGP (aggregated into a single value) against performance at different noise levels across networks. We found significant and strongest negative correlation (indicating better performance for networks with stronger anti-generalization) between CCGP and performance in network layers 2 and 3, indicating CCGP at both points is important for robust performance.

We next tested whether this way of implementing the mismatch operation could detect whether subjects made an error on single trials. As we predicted, representational alignment was significantly greater for held out correct versus error trials (0.447 vs 0.274; p < 0.01, rank sum test; Cohen’s D = 0.79). Using a signal-detection approach, representational alignment significantly discriminated correct versus error trials with an AUC of 0.59 and d’ of 0.37 (vs. chance of 0.5, p < 0.01, rank sum test). Thus, the representational alignment between a given trial’s projections into the response and color subspaces was indicative of whether subjects made an error or not. Altogether, our empirical results suggest that conflict signals in preSMA regulate the activity of forward models in preSMA, which encode color and word at the population level. preSMA also represents the actual response and a population-level version of mismatch detection can predict whether subjects made an error or not (Fig 5I).

### Error-monitoring artificial neural networks exhibit signatures of preSMA representational geometry

ANNs are typically trained to perform a given task as best as possible—in the present context, this would be to select the correct response given stimulus and task information ^51^. In contrast, here we also trained ANNs to monitor response signals and determine whether another network made an error (see also ^52^) in order to assess what representations emerge for purposes of performance monitoring. All networks received task, font color, and word meaning as input. Action Selection (AS) networks were trained to output the correct response given stimulus and task information (e.g. “red” for stimulus GREEN in the color task). Error Monitoring (EM) networks received additional input reflecting the response made by another network (or human subject) and were trained to output whether that response was correct or incorrect.

Well performing EM networks that received response input *after* stimulus and task input (schematic in Fig 5A) consistently developed representational geometries like those we identified in human preSMA. Specifically, those networks consistently exhibited color-word anti-generalization and high conflict CCGP (Fig 5B, S13). In contrast, other networks either failed to produce color-word anti-generalization, high conflict CCGP, or both (Fig 5B, S14). This included all error-monitoring networks in which response information is given simultaneously with color, word, and task information, indicating that the delayed delivery of response information (as is necessarily the case in real brains) was crucial for the emergence of this neural geometry, perhaps allowing more optimal integration of the response information with stimulus and task information).

Also crucial for the emergence of the neural geometry was the relative frequency of the types of errors made: if errors were more frequently the meaning of the word (the distractor, which is the case in human data, see Fig 1D), the networks exhibited a neural geometry with color-word anti-generalization. In contrast, if errors were instead more frequently the third, uncued option (e.g. “red” response for stimulus BLUE), networks instead exhibit above-chance color-word CCGP (Fig 5C), suggesting it is specifically the statistical relationship between the distractor and the propensity to make a corresponding error that leads to the anti-generalization aspect of the neural geometry seen in the human brain.

We next investigated how the color-word geometry exhibited by error monitoring networks contributed to error monitoring performance. We hypothesized that networks that exhibited stronger color-word anti-generalization would be more robust to increased input noise levels, due to more structured color and word encoding. Indeed, we found that networks with greater color-word anti-generalization exhibited more robust error detection performance as we added increasing gaussian noise to the network inputs (Fig 5C). Correlations between CCGP and performance were significant for networks at both layers 2 and 3, which reflect network activity immediately before and after delivery of response information respectively (Fig 5D). Interestingly, correlations were strongest for layer 2, prior to delivery of response information, suggesting the geometry plays an important role as an *error signal precursor*.

## Discussion

Humans and other animals can rapidly detect action errors even without external feedback. Internally-generated error signals in medial frontal cortex (MFC) are related to both error correction ^2,5,6,53^ and adaptation on subsequent trials ^16,20,54^, but how those signals are computed remains unclear. Here, we provide evidence that MFC encodes all signals required to detect errors by comparing a target response representation with that of the actual response, supporting the longstanding ‘mismatch’ theory of error detection ^17,29–31^. Specifically, we found that the joint activity of single neurons in preSMA, but not the dorsal anterior cingulate (dACC), encoded target (color) and distractor (word) response representations in the presence of conflicting action options during a Stroop task. Additionally, we found that preSMA encoded the actual response around the time of movement, and that population-level representations within preSMA could serve as the basis for mismatch-based error detection. These results contrast prior theorizing that error-related activity in MFC reflects reward prediction error signals delivered to dACC from midbrain regions ^33,34^, amplified response conflict on error trials ^36,37^, or a more general surprise or prediction error signal ^35,55^. Instead, our results indicate that preSMA encodes all signals necessary for the rapid detection of action errors through mismatch computations.

Our results support the hypothesis that preSMA is functionally involved in error monitoring with three main lines of evidence. The first is that the color and word signals we identified on correct trials were intact with the same representational geometry on error trials. This indicates the signals are not critical for implementing control or action selection, because error trials are precisely the times when those processes break down. Second, we found that the decodability of color and word information in preSMA, and their representational structure, was relatively stronger in sessions with greater post-error adjustments. This result is consistent with color and word representations informing error monitoring. Finally, we found that artificial feedforward neural networks that were trained to perform error monitoring exhibited similar representations of color, word, and conflict as we identified in the human data as compared to the networks trained to perform action selection. Taken together, these results suggest that preSMA internally detects errors originating from failures of control during Stroop task performance, in which control is needed to resolve the conflict arising between reading and color naming. Whether our results extend to other tasks or to broader contexts—including non-motoric cognitive errors and tasks that don’t involve a conflict between reading and another modality—remains unclear.

To perform mismatch computations, representations of the different predicted outcomes (motor actions) when control is successful must be present. One theory proposes that MFC utilizes forward predictive models—a concept borrowed from the motor control literature ^32^—to predict what response will be given if control is successful ^29^. If control is not needed, on the other hand, no prediction will be present. This hypothesis is supported by our finding that color and word decoders trained on conflict trials do not generalize to nonconflict trials. In fact, we found that preSMA activity did not differentiate between different types of nonconflict trials at all, precluding interpretations that decoders’ inability to generalize to nonconflict trials results from a simple rotation or other transformation of stimulus features in neural state-space. Instead, our results show that the forward model-derived signals in preSMA are only present in conflict trials, that is, they were conflict-dependent. This interpretation is supported by our finding that preSMA conflict signals emerged significantly earlier than color and word signals and nonconflict trials are not significantly decodeable from each other. Several studies have found that conflict signals regulate processing of task-relevant information or control parameters ^36,46–49^, including error signals ^56,57^. Here, we now show a reason for this relationship: conflict regulates error processing. Our results suggest that conflict does so by regulating the activity of preSMA forward models.

The forward model outputs were intertwined with each other—rather than being carried by distinct populations of neurons, single neurons exhibited prominent nonlinear mixed selectivity ^38,39^ for stimulus color and word, particularly in preSMA. Despite this nonlinear mixing at the single-neuron level, color and word information were linearly decodable at the population level ^39,51^. Additionally, color and word decoders exhibited anti-generalization: when applied to the opposite context, a color or word decoder exhibited below-chance performance. This result shows that color and word encoding were not independent of each other, as one would expect from representations that compete with each other. This result is incompatible with the classic conflict monitoring model of the Stroop task ^36,42^, which predicts at- or above-chance generalization depending on which part of the model is being considered. However, the addition of cross-inhibitory connections to the conflict monitoring model, reflecting competition between analogous color and word representations, led to the emergence of color-word anti-generalization in simulated data. This adjusted model therefore offers a specific circuit-level prediction for how the forward models might be implemented in preSMA. Furthermore, we found that error-monitoring ANNs with stronger anti-generalization exhibited more robust performance in the presence of increased input noise, suggesting the representational structure reflected by this anti-generalization is important for optimal error detection.

Conflict signals in preSMA were abstract despite the additional encoding of color and word information, possibly because the individual neurons that encoded conflict were largely distinct from those that encoded color and word information. This corroborates recent work by Xiao and colleagues ^44^ who found that within a given control task, intracranial high-gamma activity encoded conflict invariantly with respect to specific stimuli and responses. In contrast, Ebitz and colleagues ^40^ recently reported that individual neurons in dACC encode conflict by amplifying task-relevant information (i.e. font color) and suppressing task-irrelevant information (i.e. word meaning; also see ^41^). We did not find that task-irrelevant information was suppressed. Rather, in preSMA, the task relevant and irrelevant information was represented equally strongly and in dACC, neither was decodable. Intriguingly, the error monitoring ANNs we analyzed also exhibited abstract conflict signals despite not being trained to detect conflict. Our results thus suggest that conflict signals may emerge specifically because of the raised likelihood of making errors on conflict trials, as opposed to an increased need for control.

Crucially, we found that the actual response that subjects enacted was encoded in preSMA. Where did this information come from? Corollary discharge theory (aka reafference or efference copy; ^58^) holds that copies of motor signals are transmitted to sensory or other cortices to diminish prediction errors related to one’s own behavior ^59–62^. In primates, corollary discharge signals are well-described for saccades ^62,63^, with much less known about the pathways providing the same information for finger and hand movements. Our results build on this literature by showing that the specific action taken by our participants was decodable in the preSMA and dACC, a signal presumably provided to these areas through similar corollary discharge signals. While the precise pathways remain unknown, we hypothesize that action selection circuits in the basal ganglia route information about the selected action to MFC through the thalamus ^22,29,64^. To our knowledge, our study is the first case where encoding of the actual and target (or intended) action was disentangled and directly demonstrated in humans. Our study thus provides evidence for corollary discharge theory’s applicability to humans.

The mismatch model of error computations suggests that representations of the target and actual response are compared to detect errors ^17,22,29,31^. We observed mixing for representations at the single-neuron level, requiring a population-level approach to the mismatch computation. Therefore, we used representational alignment as a metric, which assesses the cosine similarity of response and color (target) representations at the population level. Error trials had lower alignment than correct trials, showing that this metric could be used to predict whether a subject made an error without relying on external feedback. This suggests that binary error detection in the brain could arise from a threshold operation on this metric. In addition, by virtue of providing a continuous value, representational alignment might also naturally provide information about the magnitude and, considering the measure’s inputs, types of errors, which could be utilized for error correction ^2,5,6^ or behavioral adjustment when feedback is insufficient ^65^.

Taken together, our results are compatible with an actor-critic framework of action selection and performance monitoring implemented in a distributed network with basal ganglia serving as the actor and MFC as the critic ^29^. In this model, basal ganglia implements action selection (the actor) and dACC tracks the action selection process to detect conflict. Conflict signals in dACC are transmitted to preSMA, thereby engaging the conflict-dependent forward models and error detection mechanisms described in our present study. Error signals are then routed back to dACC which adjusts control parameters, such as response thresholds, for the following trial, leading to post-error slowing. In this framework, psychiatric illnesses with implicated error monitoring including OCD ^8,9,11,13,15^, may involve preSMA dysfunction (see also ^66^). Notably, current surgical and stimulation approaches to OCD treatment focus on the ACC ^67,68^, but our results suggest that the preSMA may be an important alternate target to consider. Altogether, our study advances our theoretical and computational understanding of preSMA’s and dACC’s diverging roles in performance monitoring and has important implications for future research and clinical applications.

## Acknowledgments

We thank Jeffrey Schall, Stefano Fusi, Kai Miller, Clayton Mosher, Natasha Kurilenko, Hristos Courellis, Amirsaman Sajad, Ben Corrigan, Pranavan Thirunavukkarasu, and Steven Errington for discussion. We thank the Cedars-Sinai epilepsy monitoring unit physicians and staff for their support and the patients for their participation when this data was originally acquired (this study is a re-analysis of existing published data). This project was funded by the NSF (BCS-2219800 to UR) and a Center for Neural Science and Medicine at Cedars-Sinai postdoctoral fellowship (to JG).

## Methods

Full details of the utilized task, patient population, and data preprocessing can be found in Fu et al., (2022). Briefly, the subjects were 34 epilepsy patients (87 sessions total) implanted with hybrid depth electrodes for seizure monitoring. Spikes were detected and sorted with a template-matching algorithm and we only analyzed well-isolated single neurons. Subjects performed a speeded version of the classic color-naming Stroop task (word & color options were both red, green, and blue). Responses were followed by a blank screen (1 s), and then feedback (“correct”, “incorrect”, “too slow”; 1 s), and stimuli sequences were randomized.

### Behavioral analysis

Differences in reaction times for conflict versus nonconflict trials and post-correct versus post-error were calculated using mixed effects models with conflict and prior trial accuracy as fixed factors. Models were fit with random intercepts according to session ID to account for variance across individuals and recording sessions. Differences in error rates for conflict versus nonconflict were calculated similarly with conflict as a fixed factor and random intercepts according to session ID. We used only correct trials with response times less than 2 seconds following stimulus onset for this analysis.

### Single-neuron Analyses

For individual neurons, we recorded response times following spike sorting (see Fu et al., 2022 for details). We calculated peri-stimulus time histograms of firing rates using a 500 millisecond window (where the leading edge of this window corresponds to the times shown in, e.g., Fig 1G). In other words, the shown firing rate at time 1.0 s relative to stimulus onset corresponds to the average firing rate in the window [0.5 1] s relative to stimulus onset.

For statistical analysis of firing rates, we used 3×3 ANOVAs implemented in Matlab R2023A (anova function) or Python (statsmodels functions ols and anova_lm). Color and word were included as the two main effects. For the joint analysis with conflict we included conflict as a covariate in the formula (e.g. firingrate ∼ color*word + conflict) and then converted regression results to F statistics using statsmodels’ function anova_lm.

For visualization of neurons’ response profiles (Fig 1J) we took every neuron’s condition-average firing rate in the stimulus window ([0.5, 1] s relative to stimulus onset) and then z-scored activity across conditions in order to represent response profiles succinctly in a manner consistent across neurons. We then ordered neurons so that the Euclidean distance between the 9×1 vector of response profiles of consecutive neurons was minimized.

### Population decoding of color and word

We formed pseudopopulations by aggregating individual neurons’ single-trial PSTHs according to the presented color and word. We used only correct trials with response times less than 2 seconds following stimulus onset for this analysis. Trials were shuffled to destroy noise correlations, which can reduce decoding accuracy ^69^. We then sub-sampled trials to have an equal number among all conflict conditions, which left us with 8 trials for each of 6 conflict conditions.

For decoding, we applied a support vector classifier to pairs of conflict conditions that matched on either color (word decoding) or word (color decoding) features. We utilized a leave-two-out stratified cross validation procedure to assess out-of-sample test accuracy because of the low trial numbers. To assess significance, we constructed a null distribution of decoding accuracies by repeating the decoding procedure on the same data with shuffled condition labels. For single-timepoint decoding (e.g. Fig 2B) we repeated this procedure across 20 random initializations of the pseudopopulation, comparing to 50 random shuffles for each initialization, for a total of 1000 bootstrapped datapoints for the null distribution. We categorized a particular timepoint’s decoding accuracy as significant if it was above the 95^th^ percentile of accuracies within this null distribution. For time-resolved decoding (e.g. Fig 2B right) we compared a single pseudopopulation against a null distribution of 1000 shuffles.

To assess cross-condition generalization accuracy, we took decoders trained on a pair of conflict stimuli that differed only on one feature (e.g. Blue/red vs Blue/green) and tested them on corresponding pairs in the opposite context. For instance, we trained a color decoder to classify Blue/red vs Blue/green and measured CCGP by testing its performance in classifying Red/blue vs Green/blue. We ensured labels used for training and testing corresponded to the desired feature. For assessing significance, we similarly trained decoders on data with shuffled labels to construct a null distribution and categorized decoding accuracy as significant if it was above the 97.5^th^ percentile or below the 2.5^th^ percentile of null accuracies (corresponding to a 2-sided test with alpha of 0.05).

### Population Decoding of Conflict

To assess the presence of conflict signals at the population level (e.g. Fig 3B) we similarly formed pseudopopulations according to whether a given trial was conflict or nonconflict (shuffling color and word information), subsampled trials for equal numbers and then performed decoding and bootstrapping to assess significance as above.

To assess the geometric structure of conflict signals we once again formed pseudopopulations according to both color and word stimulus features, for a total of nine conditions. We measured CCGP by training conflict decoders on all data leaving out one pair of conflict and nonconflict conditions, and then testing those decoders on data from the left out conditions. We repeated this analysis a total of 18 times (once for every possible pair of conflict and nonconflict conditions) and report the average (plus individual traces) in Figure 3C. We used logistic regression for this decoding analysis to reduce fitting time to reduce the relatively large number of condition combinations.

### Pairwise Decoding and Latency Analysis

We extended our color and word decoding analysis (using the same pseudopopulation and inclusion criteria) to all condition pairs. We once again used leave-two-out stratified cross-validation with support vector classifiers to assess decoding accuracy. We grouped pairs of conditions into three types: conflict-conflict pairs (two conflict conditions), conflict-nonconflict pairs (one conflict condition and one nonconflict condition; note that this corresponds to conflict signals), and nonconflict-nonconflict pairs. Conflict-conflict condition pairs could be further subdivided into pairs that shared the same word but differed in color or vice-versa, respectively corresponding to the color and word decoding we analyzed earlier, pairs that shared no stimulus features (e.g. Red/blue and Blue/green), and pairs that were inverted in terms of color and word features (e.g. Red/blue and Blue/red).

To compare the latency of these different signals, we took as latency the first timepoint greater than 400 ms post-stimulus onset that reached significance compared to the null distribution (training on data with shuffled labels; 1000 bootstraps), and compared groups using an independent samples t-test. The difference between groups remained significant using a threshold metric (e.g. first timepoint above 70% decoding accuracy) at a variety of threshold levels, and did not depend on the inclusion of all types of conflict-conflict condition pairs versus just color-word decoding pairs.

### Demixed PCA

We utilized demixed principal components analysis (dPCA) to visualize the dynamics of preSMA activity following stimulus onset. We took the average activity across time for every neuron in each of the 9 conditions to form a condition-average pseudopopulation and utilized dPCA (implemented through the Matlab toolbox (REF)) to assess the relative contribution of color, word, color-word interactions, and time on activity following stimulus onset. We projected activity onto the first two color-word interaction components and subtracted a pre-stimulus baseline (average projected activity for each condition in the period [-0.5, 0] s relative to stimulus onset) to assess how stimulus information affects dynamics in preSMA.

### Generalization to Error Trials

Due to low error rates in most patients, error trials were relatively infrequent in our dataset, with some patients even performing zero trials incorrectly. Therefore, it was not feasible to perform a full decoding and CCGP analysis on error trials. Instead, we formed pseudopopulations from correct trials grouping data according to color and word features, restricting our sessions to those with at least 6 errors on conflict trials. Then, we formed a matched pseudopopulation from exclusively error trials without replacement, imputing missing data with NaNs. To assess whether decoders trained on correct trials would generalize to error trials, we fit logistic regression classifiers to correct trials and applied them to error trials, ignoring neurons with missing data. Logistic regression models the log-odds that a given sample comes from one class versus another by fitting the regression equation:

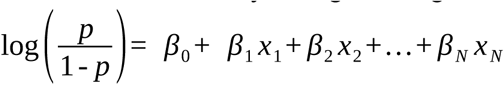

Where *p* is the probability that a given pseudotrial belongs to class 1 (wlog), *x_i_* is the firing rate of neuron *i* in that pseudotrial, and N is the number of neurons in the pseudopopulation. As a linear equation without interaction terms, the *β_i_* coefficients reflect the relative effect of each neuron’s firing rate independent of all other neurons. So, for generalization we restricted the logistic regression to the neurons that actually had data on a given error pseudotrial, taking the predicted class output of the restricted logistic regression as our prediction. Then, to test the significance of this analysis we trained decoders on the same correct trial pseudopopulations with shuffled labels and then applied them to the properly labeled error trial pseudopopulations. We repeated this shuffling procedure 1000 times and determined that generalization to error trials was significant if it was above the 95^th^ percentile of the bootstrapped distribution.

### Post-Error-Slowing Comparative Analysis

We sessions into two groups based on whether they exhibited above- or below-median degrees of normalized post-error slowing. As described in Fu et al., (2019), we calculated post-error slowing as the difference in reaction time between the trial before and after an error (matching for conflict or nonconflict), divisively normalized by the session-average reaction time on correct trials. If a session did not have at least 5 conflict-matched pairs of trials surrounding an error trial, we omitted it from this analysis. This left us with 48 sessions with a total of 359 neurons in preSMA (195 from high PES sessions, 164 from low PES sessions).

We next sub-selected 150 neurons from each of these two session groups to form pseudopopulations, on which we performed the binary color and word decoding & CCGP analysis as described above (e.g. Figure 2A, 2D). We then repeated the analysis with *neurons* randomly partitioned into two groups (as opposed to partitioned into groups according to PES score) to calculate a ‘null difference’ between groups. To determine whether the difference between high- and low-PES groups was significant, we repeated this pseudopopulation formation and decoding/CCGP procedure 1000 times, forming distributions for the real difference between groups and null difference between groups. Then we performed a Wilcoxon sign-rank test (implemented in scipy) on these distributions to assess significance.

### Marginal Decoding of Color, Word, and Actual Response

Correct trials are, by definition, trials where the color and actual response are equivalent. Therefore, to assess whether actual response is represented alongside color and word information, we analyzed correct and error trials together. We formed *marginal* pseudopopulations that were labeled according to the color, word, or actual response (hereon referred to as *features*), while randomly shuffling the other two features. We first calculated how many error trials were available for a neuron when a feature was red, green, or blue (hereon referred to as factors). We formed pseudopopulations from a mixture of these error trials and correct trials such that each factor had a number of trials equal to the total number of errors. For instance, if a neuron had 5 error trials where ‘red’ was pressed, 4 error trials where ‘green was pressed, and 3 error trials where ‘blue’ was pressed, totalling 12 error trials, we added 7 correct trials where ‘red’ was pressed, 8 correct trials where ‘green’ was pressed, and 9 correct trials where ‘blue’ was pressed, for a total of 36 trials. We then randomly selected 10 trials for each factor from each neuron to form pseudopopulations. These were randomly shuffled within factors before activity was concatenated to ensure that a given pseudotrial may contain, for two neurons recorded simultaneously, a mixture of information from correct and error trials. If a given neuron did not have 10 trials available for each factor it was not included in the pseudopopulation.

We used support vector classification to decoding color, word, or response from the marginal pseudopopulations, utilizing stratified leave-three-out cross validation to assess accuracy while keeping classes balanced. Then we shuffled training labels to assess chance level accuracy. Due to the possibility of being biased by the small number of error trials, we repeated this pseudopopulation construction and decoding procedure 1000 times, randomly mixing trials across neurons for each pseudopopulation. For each bootstrap we retained one ‘true’ decoding accuracy value, and one ‘null’ decoding accuracy value, and then used an independent samples t-test on the final obtained distributions to ascertain whether decoding accuracy was above-chance. We repeated this procedure for a variety of trial cutoff numbers (balancing the tradeoff between number of trials and number of neurons available for decoding) and for a similar procedure where we used fewer correct trials/relatively more error trials and found largely the same results.

### Representational Alignment Procedure

To assess whether our decoding results for color and response were sufficient to compute an error signal, we formed pseudopopulations as in the section above from a mixture of error and correct trials. We held out a single correct and error trial for the same stimulus where response matched the color and word respectively. We then fit support vector classifiers to the data using a one-versus-rest procedure (LinearSVC from sci-kit learn), and extracted the coefficients of the fitted classifiers to define a “red”, “green” and “blue” coding vector separately for the color and response pseudopopulations. We then projected the activity from the held-out trials onto each of these vectors, effectively projecting activity into a color or response subspace, from which we defined projection vectors e.g. 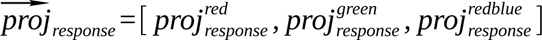. We defined representational alignment as the cosine similarity of the color and response projection vectors, i.e. 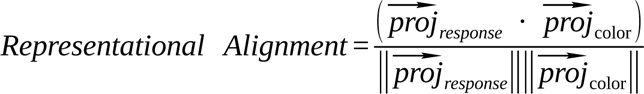. Based on the reasoning that corresponding indices within the color and response projection vectors should both be high or low for correct trials (i.e. the projection vectors point in similar directions) due to the response and color matching, whereas projection vectors should be pointing in different directions in error trials, we reasoned that representational alignment should be high for correct and low for error trials.

We calculated representational alignment by randomly holding out a correct and error pseudotrial, and fitting decoders on remaining data. We repeated this procedure 1000 times for each condition (total 6000 trials), resampling the held out pseudotrials each time. To assess the utility of this measure for predicting errors we calculated the AUC and d’ of single-trial representational alignment values with 0 (error) and 1 (correct) as labels. We also repeated this procedure on null data (maintaining correct/error held-out trials but shuffling color/response labels when training) and on properly-labeled data from a baseline period and found that the difference between correct and error trials in representational alignment, as well as AUC and d’ was significantly greater in properly labeled data compared to null or baseline valued (all p < 0.01, rank sum test).

### Computational Modeling: Conflict Monitoring Model & Cross-Inhibition Model

We based our computational models of the Stroop task on the circuit model presented by Botvinick and colleagues (2001). We implemented this circuit as a continuous rate model that consisted of 3 color, 3 word, 3 response, two task, and one conflict node connected hierarchically so that input to the color, word, and task nodes drives selection of the ‘correct’ response. Nodes of the same type additionally had mutual inhibitory connections. Activity was governed by the equations:

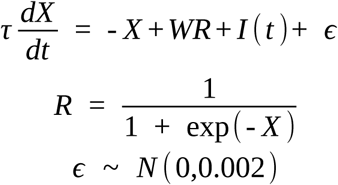

Where *τ* = 250 *ms* is the synaptic time decay constant, *X ɛ* ℝ^12^ *^x^*^1^ is the vector of synaptic current variables for each of the 12 nodes, *R ɛ* ℝ^12^ *^x^* ^1^ is the activity state, *W ɛ* ℝ^12^ *^x^*^12^ is the connectivity matrix which implements the model given in Figure 2, *I* (*t*) *ɛ* ℝ^12^ *^x^* ^1^ is the time-varying vector of input currents that encode the stimulus on a given trial, and *ɛ* is normally-distributed white noise that is added at every timepoint.

Activity was simulated at a resolution of 1 millisecond using the Forward Euler method. Task input was given immediately, and stimulus input was given after 1 second had elapsed. This was the point used for ‘stimulus onset’ in further analyses. We simulated 4 seconds total of activity for 500 trials, randomly selecting the stimuli with equal probability. Reaction time and whether the model performed correctly was determined by the time at which a response node first passed an activity level of 0.75. For the decoding analyses we confirmed more than 300 correct conflict trials and then proceeded as with the human data. We used four-fold cross-validated logistic regression to investigate color and word representations owing to the high number of simulated trials.

Network parameters for connections between distinct node types in the conflict monitoring model were as follows: task → color/word = 3, color/word → response = 2, response → conflict = 0.2, and conflict → task = 0.2. Nodes of the same type were also connected with inhibitory projections to stabilize activity: task → task = −2, color → color = −3, word → word = −3, response = −3. Response nodes also experienced self-excitation (strength 3) as is common in winner-take-all models.

The cross-inhibition model utilized largely the same parameters with the addition of cross-inhibitory connections between color and word nodes corresponding to the same feature (e.g. red color and red word mutually inhibited; strength −3). To account for the additional inhibition in this model we reduced the strength of lateral inhibition connections among color and word nodes from −3 to −1, which led to similar response times as the conflict monitoring model (see Fig S6)

### Artificial Neural Networks

#### Network Architectures

We trained and analyzed a variety of artificial neural networks in the context of our task. Networks could be broadly broken down into two categories based on their objective. Action Selection (AS) networks received as input the stimulus color and word (both red, green, or blue), as well as the task (color-naming or word-naming task), and had to output the appropriate response (red, green, or blue) through combining that information. In contrast, Error Monitoring (EM) networks received the same inputs in addition to a given ‘response’ (red, green, or blue) and had to determine whether the response was correct or incorrect based on all inputs.

The networks were further varied by when (i.e. to which layer) they received the various inputs. We trained AS and EM networks that received all inputs to the first layer of the network, AS networks that received either the stimulus color and word information at layer 2 or 3, AS networks that received task information at layer 3, and EM networks that received response information at layer 3.

For comparison across all tasks we used 3 hidden layers of 50 neurons each. Hidden layers used ReLU activation and the final output layer used softmax activation. Our results were consistent with different number of layers, layer sizes, and activation functions.

#### Network Inputs

Inputs were one-hot-encoded and concatenated to form one or more one-dimensional vectors that were input to the appropriate network layers. For instance, the EM receiving input corresponding to the word “red” written in blue ink in the color task and had to determine whether the response red was incorrect received the 8×1 vector [0 0 1 1 0 0 1 0] as input to the first layer (the first three indices corresponding to the color, then the word, then the task) and the 3×1 vector [1 0 0] as input to the third layer in addition to the output of layer two. Finally, to simulate biological input which is generally noisy we added a small amount of gaussian white noise (STDEV 0.05) to each of these inputs prior to training.

For AS networks individual trials (i.e. stimulus/task combinations) were randomly selected with uniform probability. For EM networks stimulus/task combinations were randomly selected with uniform probability, and the input response was equally split between the color and word for that trial. This resulted in a 0% error rate on nonconflict trials and 50% error rate on error trials, with errors always being equivalent to the distractor stimulus. We used this protocol to more evenly balance the training data.

#### Network Training

We trained 50 networks for each architecture described above (each initialized randomly) using an ADAM optimizer with cross-entropy loss, where network outputs were compared to a one-hot encoded output (e.g. [1 0 0] for desired output ‘red’ for the AS networks, [0 1] for desired output ‘error’ for the EM networks). We trained for 5000 epochs with a batch size of 10 trials per epoch, updating network weights after each epoch. Every network reached above 95% accuracy in performing their respective objectives.

#### Decoding Color and Word

After training, we simulated network activity for 2000 randomly generated trials using the same parameters as in the training data. We omitted from further analysis those trials in which the network gave an incorrect output or, in the case of the EM networks, where the inputted response was incorrect. We separately applied the same color-word decoding & CCGP protocol as we did to the human data to the activity at each of the three network layers, with the modifications that we used logistic regression classifiers with four-fold cross-validation for training, and area-under-the-curve as our accuracy metric to reduce computation time. For a reduced dataset the findings were similar using the same decoders/metrics as human data. We also applied the same conflict generalization analysis as we applied to human data. Decoding accuracies for color, word, and conflict were unvaryingly at 100%.

#### Types of Error Analysis

We trained EM networks with response input at layer 3 with different ratios of types of errors. Specifically, we defined the probability that a given error on conflict trials would match the distractor (i.e. response ‘red’ for “red” written in blue ink) versus match the third, uncued option (i.e. response “green” for “red” written in blue ink). We parameterized this as the probability that an error matches the distractor, ranging from 0-100% (note the probability of the error matching the third option is 1 minus this probability). We trained 20 networks with the same random weight initializations using input data where this probability ranged from 0 to 100% in incremennts of 10% and then applied the same decoding procedure to the networks as described above.

#### Robustness-to-Noise Analysis

We took the 50 trained EM networks with response input at layer 3 and tested their capacity to perform error detection in the presence of increasing input noise. We simulated trials with the same parameters used for training except the standard deviation for white noise (‘noise level’) varied between 0 and 2 (note it was 0.05 in the original training dataset). We simulated 5000 trials for 101 equally spaced noise levels between 0 and 2 for each of the 50 networks and then tested whether the networks outputted the correct output (error or correct) given the inputs. Network output was defined as which output neuron was more active (a fairly lax threshold).

We next correlated network performance at each of these noise levels against the layer-specific color-word CCGP exhibited in the prior analysis (i.e. for noise level 0.05; we averaged color->word and word->color CCGP for this analysis). We fit linear models using the linregress SciPy function and extracted correlation coefficient and p-values from the fitted models. Fitting failed for noise levels between 0 and 0.08 because under these conditions all networks exhibited perfect performance.

