## Supplementary Figures for "Autonomous error detection is enabled by conflict-dependent forward models in human medial frontal cortex"

### Supplementary Information

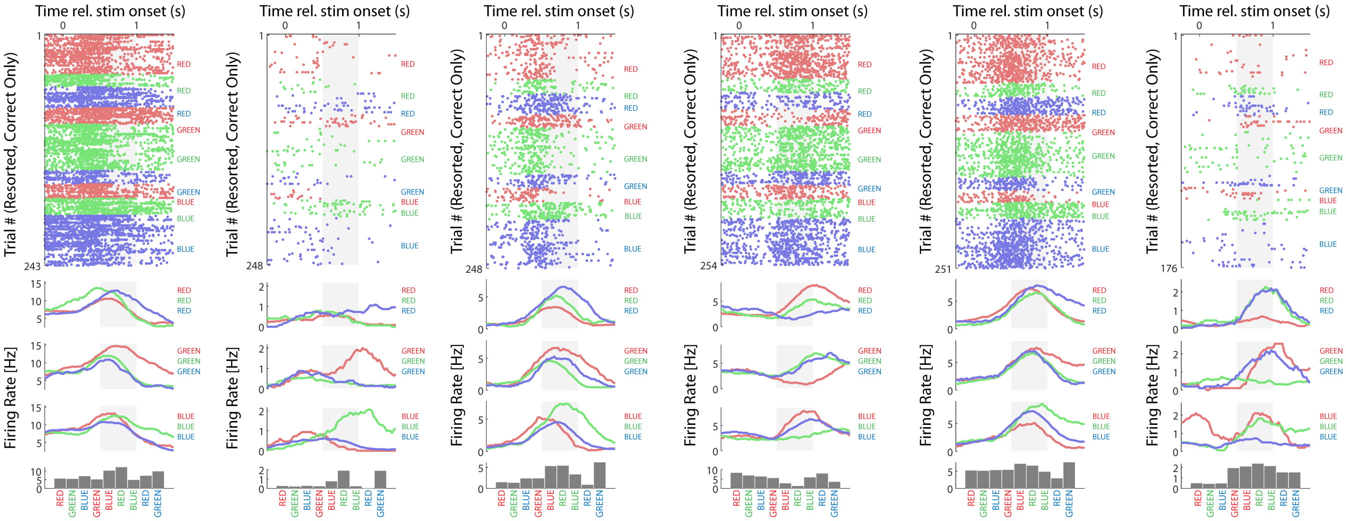


**Fig S1: Individual Neuron Examples.** Rasters (top), PSTHs (middle), and average FR in the window [0.5,1] (bottom) are as described in Fig 1.

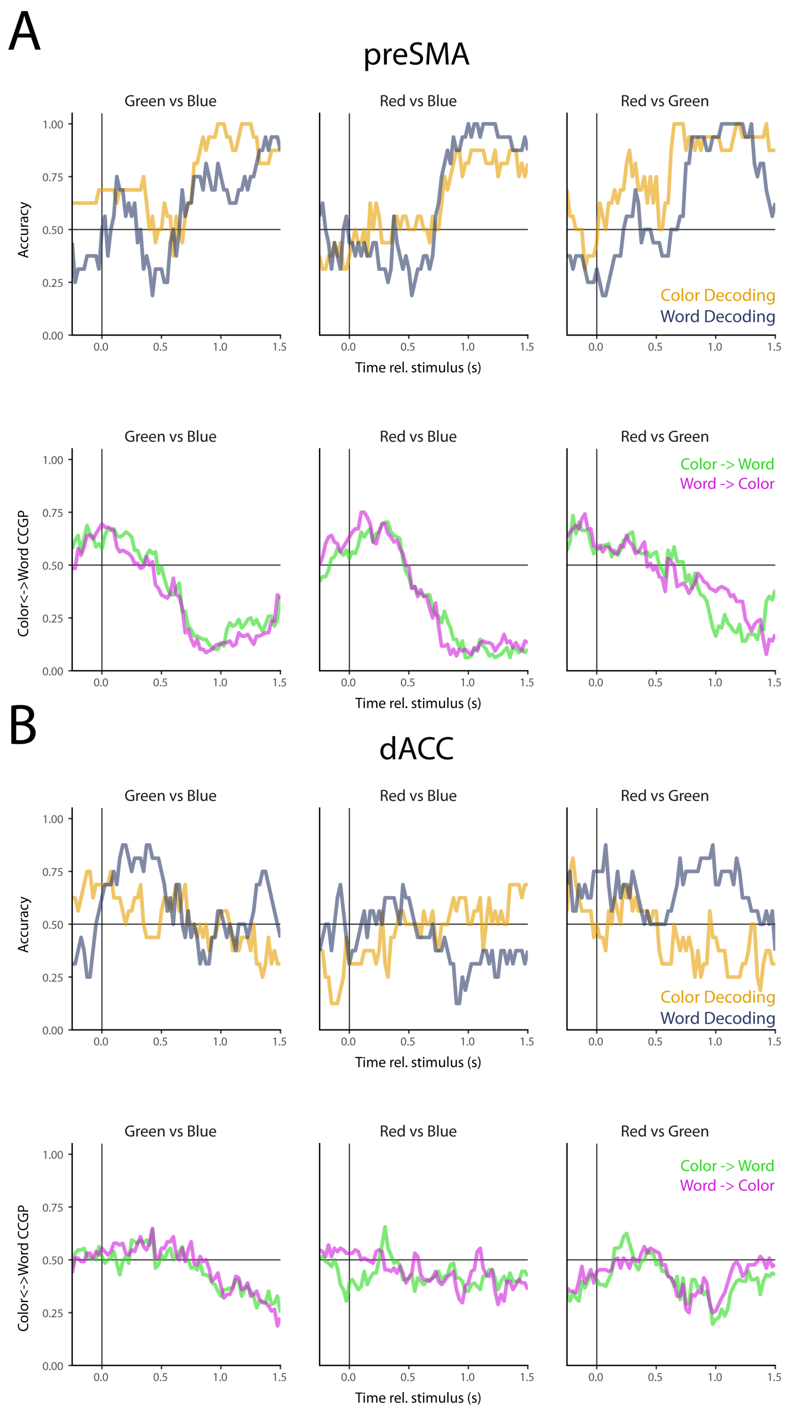


**Fig S2: Color and word decoding in preSMA and dACC.** We show the average pairwise decoding accuracy & CCGP for individual decoding pairs (see Fig 2A, 2E).

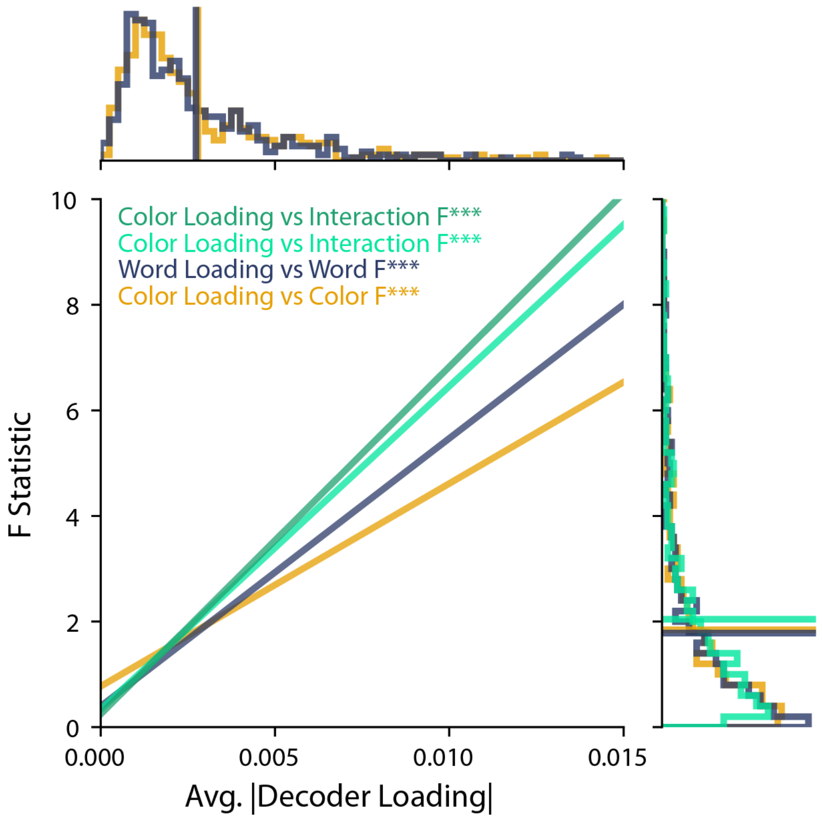


**Fig S3: Correlations between color/word decoder loadings and F statistics.** We took the absolute value of the color and word decoder loadings (averaged across all three pairwise options) and correlated them against single-neuron F statistics for Color, Word, and Interaction. Decoder loadings were more strongly correlated with interaction F statistics than with main effect F statistics. Top and right are histograms of F statistics and decoder loadings.

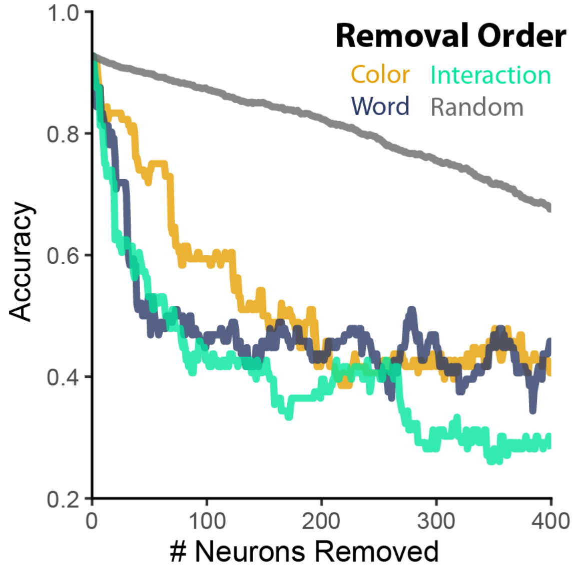


**Fig S4: Virtual Ablation Analysis.** We performed a virtual ablation analysis to assess the role of neurons that tuned for different stimulus features in supporting our decoding results within preSMA. We performed the same decoding analysis as in Fig 2B (restricted to the period [0.5, 1] relative to stimulus onset) while iteratively removing the neurons with the largest F statistics (or randomly). Decoding dropped off sharply when removing neurons according to their F statistics compared to randomly, indicating our population decoding results are driven by neurons with strong single-neuron tuning.

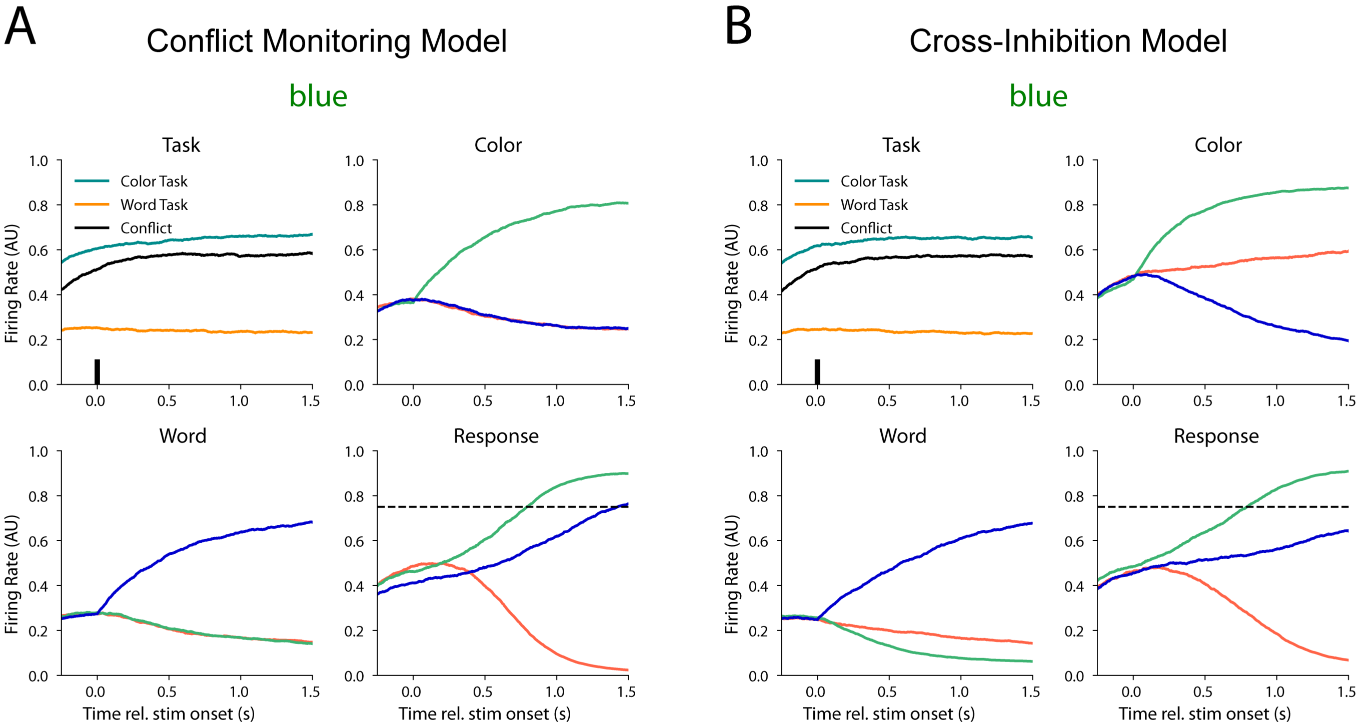


**Fig S5: Example of simulated activity**. Shown are individual simulated trials for the Conflict Monitoring (A) and Cross-Inhibition (B) models (model schematics in Fig 2E and 2I respectively). Model response time was determined by the time at which activity passed a threshold of 0.75, and model response was the unit that first reached that value. Competition in the models is driven by different input to the response layer from the color and word units to the response units. In the Cross-Inhibition model, inhibitory connections between analogous color and word units lead to relative suppression effects. For instance, the blue color unit shows a lower firing rate than the red color unit, because the blue color unit is being inhibited by the blue word unit, which is activated by the distractor input. This relative suppression is absent in the Conflict Monitoring model, and underlies the color/word anti-generalization seen in this model.

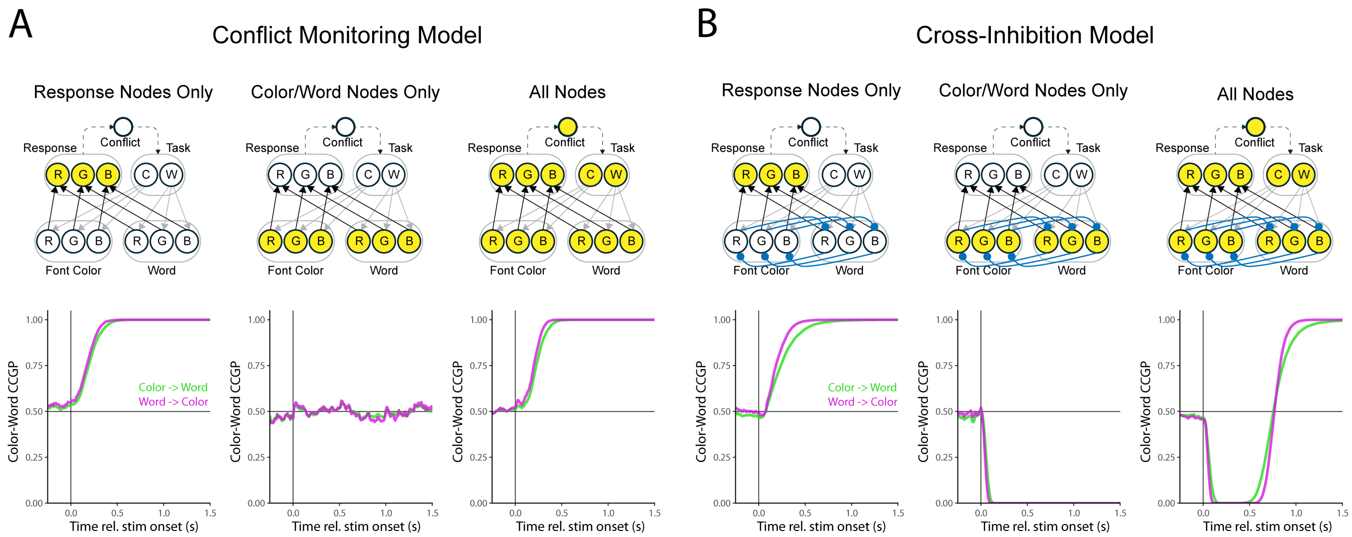


**Fig S6: Decoding activity from model sub-circuits.** We applied the same decoding analysis in Fig 2F & 2J to sub-circuits within the Conflict Monitoring (A) and Cross-Inhibition (B) models. The nodes whose activity were used for decoding are colored yellow in the model schematics above. Note that color and word decoding was always at ceiling, so we restrict our investigation to CCGP. In the conflict monitoring model, above-chance CCGP is driven by the response units: when restricting to the color and word units, CCGP is at chance. In the cross-inhibition model, CCGP for the response units is above-chance, whereas CCGP in the color and word units is below chance. When considering the whole network, CCGP first trends below- and then increases to above-chance. This reflects earlier emergence of information in the color and word units compared to the response units.

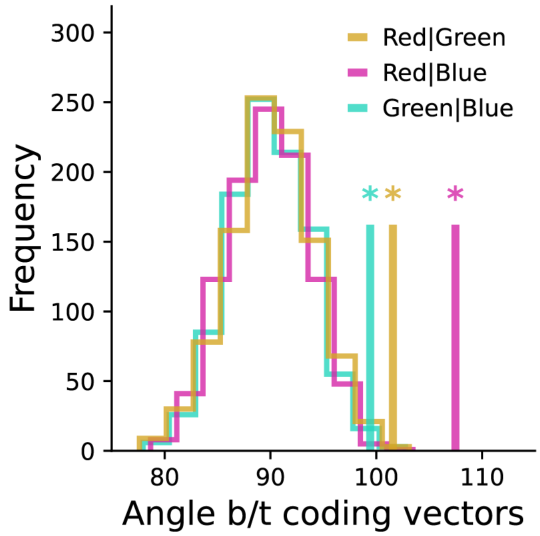


**Fig S7: Angles between corresponding color and word decoders.** We compared the angles between color and word decoders for corresponding pairs of stimuli. We find that the angles are all significantly greater than chance. Vertical lines are empirical angles between decoders, histogram is the null distribution (1000 bootstraps, results obtained by training on shuffled labels).

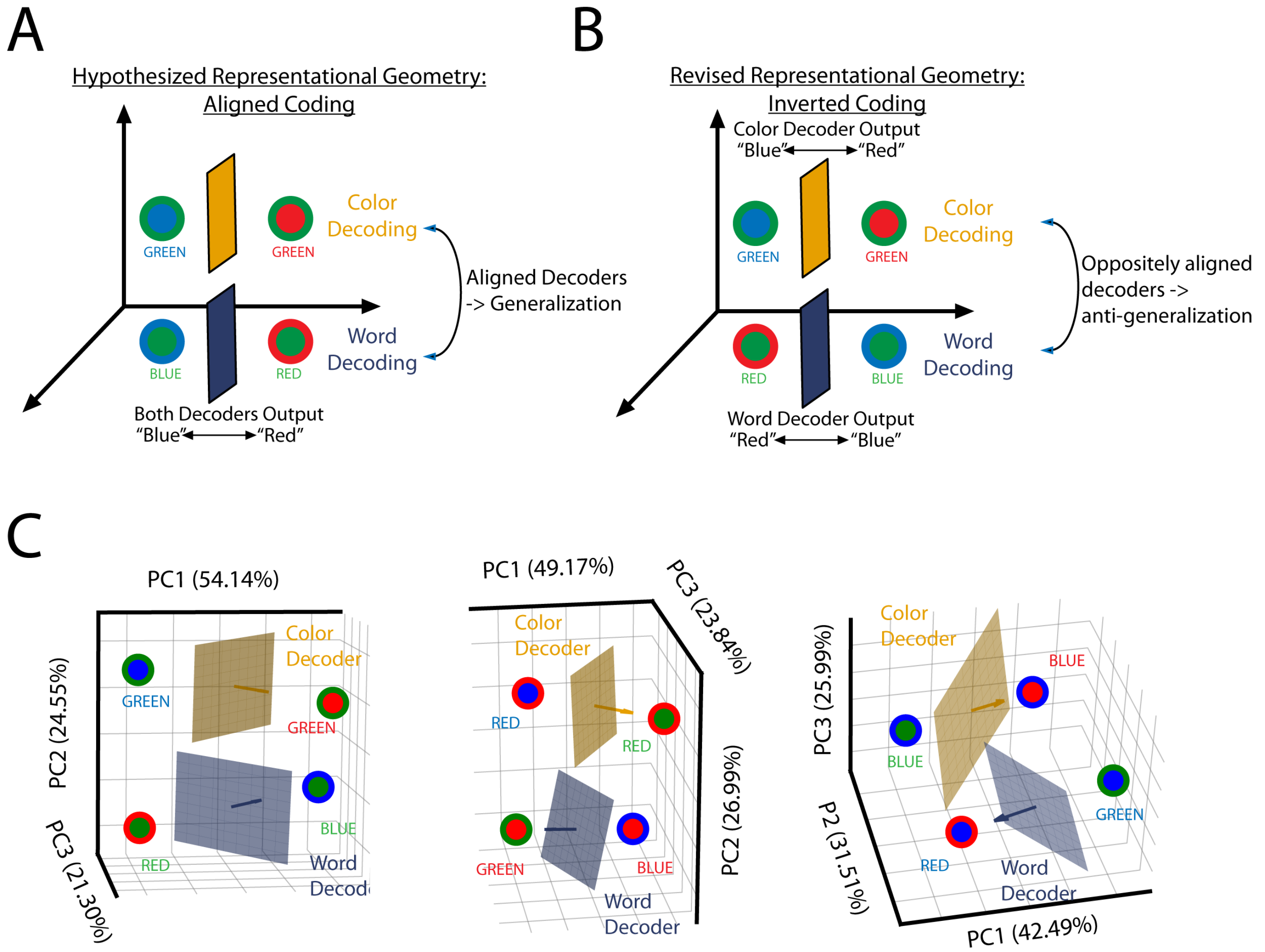


**Fig S8: Schematic for anti-generalization and visualization of preSMA activity. (A-B)** Schematics for how anti-generalization can emerge: if color and word information is organized in the same direction in state-space as in (A), decoders should exhibit generalization. On the other hand, anti-generalization should reflect color and word information being organized in opposite directions. **(C)** Principal component visualization of preSMA activity. We averaged activity across trials for each neuron in the 6 conflict conditions, and then visualized the 3 sets of 2 stimulus pairs that correspond to our color-word decoding and CCGP results. Schematic color and word decoding planes are shown for illustrative purposes. The organization of activity along the firs three principal components corresponds to the inverted coding scheme described in panel B.

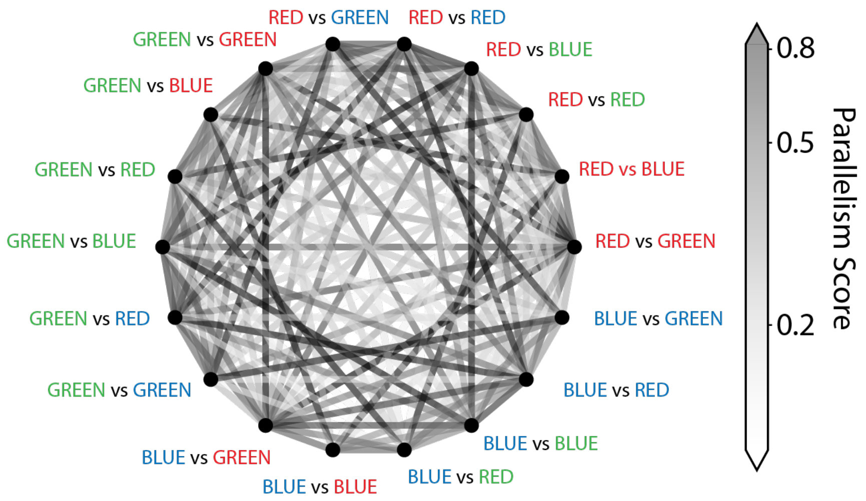


**Fig S9: Parallelism Scores for Conflict Decoders.** We found the conflict coding vectors obtained from decoders trained on pairs of conditions (18 possible condition pairs; coding vectors obtained from logistic regression models fit to data from individual pairs). We then calculated the parallelism score between every pair of coding vectors as the dot product of the two normalized vectors (306 possible coding vector pairs). The color of lines between condition pairs in the panel reflects the parallelism score between those two coding vectors. We assessed the significance of each parallelism score by comparing the value to the 95$^{th}$ percentile of a null distribution where conflict decoders were trained on data with shuffled labels from the same condition pair (1000 bootstraps). Of 306 coding vector pairs, only 5 did not have significantly above-chance parallelism scores.

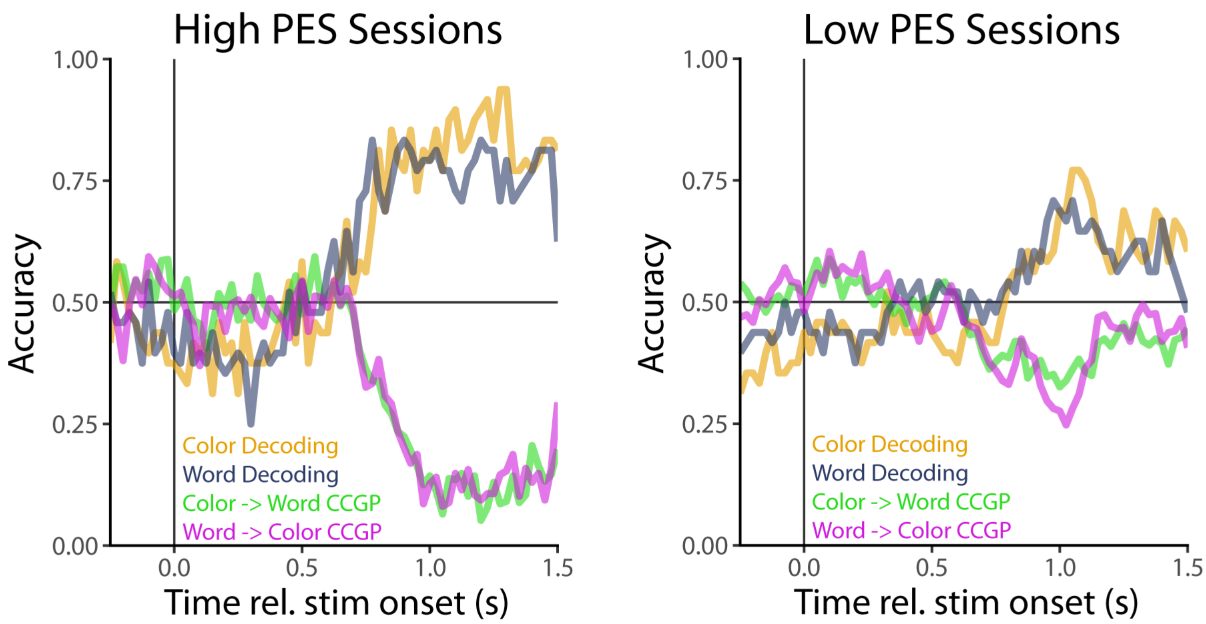


**Fig S10: Decoding Accuracy over time for high/low PES groups in preSMA.** We conducted the same time-resolved decoding procedure as in Fig 2B separately for neurons recorded during sessions with high or low PES scores. As in the case for the single-timepoint, high PES sessions have far stronger and more structured encoding of color and word information.

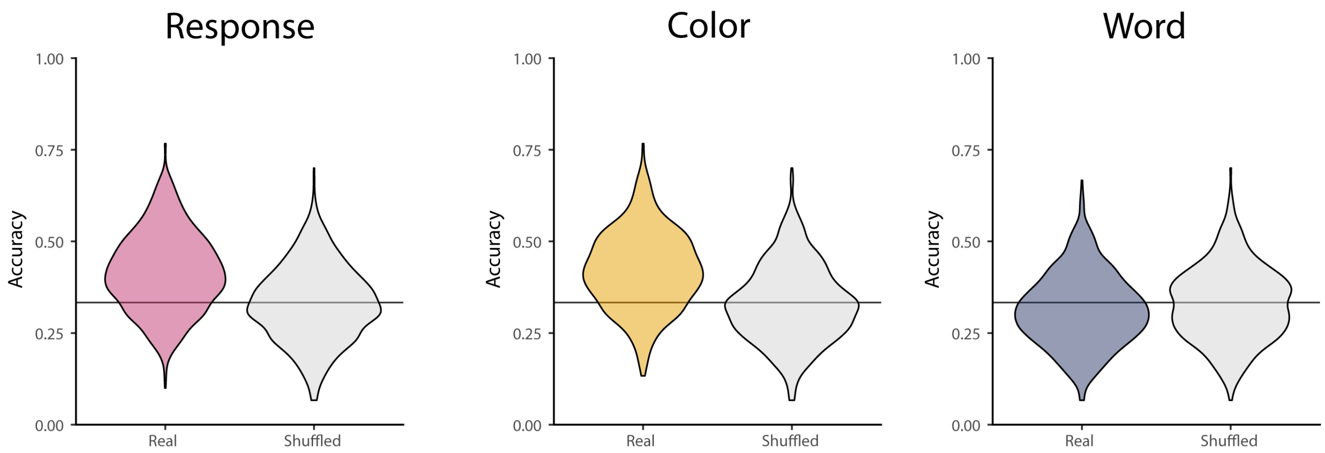


**Fig S11: Decoding Distributions for marginal decoding of response, color, and word.** Shown are distributions of decoding accuracies obtained from properly-labeled pseudopopulations (left in each panel) and randomly-labeled pseudopopulations (right in each panel) in the period 500 ms around button-press. Decoding accuracies were highly variable due to the infrequency of error trials (see methods), but response and color decoding was elevated compared to the shuffled distribution in this period.

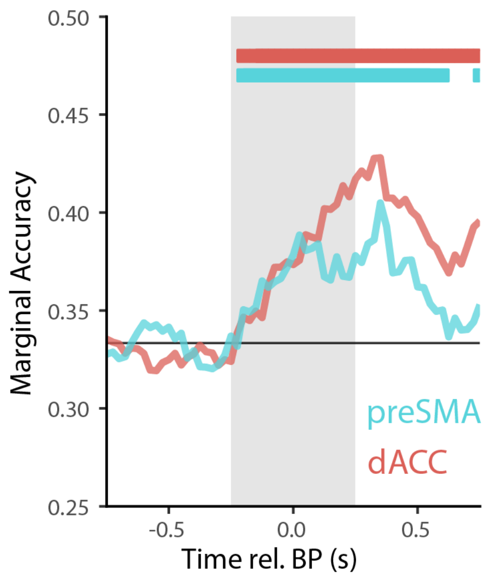


**Fig S12: Decoding response in preSMA and dACC.** We decoded actual response from preSMA and dACC and found that we could decode response from both regions with nearly the same time-course. Horizontal bars are periods of significance compared to a null distribution (1000 bootstraps; FDR-corrected).

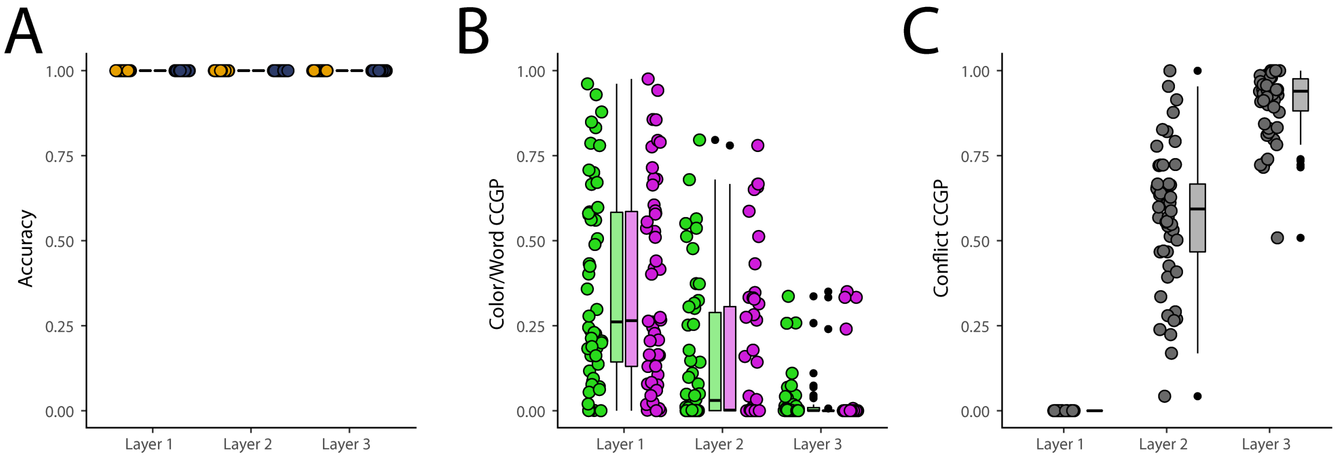


**Fig S13: EM network decoding results.** Accuracy (A), color/word CCGP (B), and conflict CCGP (C) for the error monitoring network shown in Figure 5A. Each point reflects results from a single trained network.

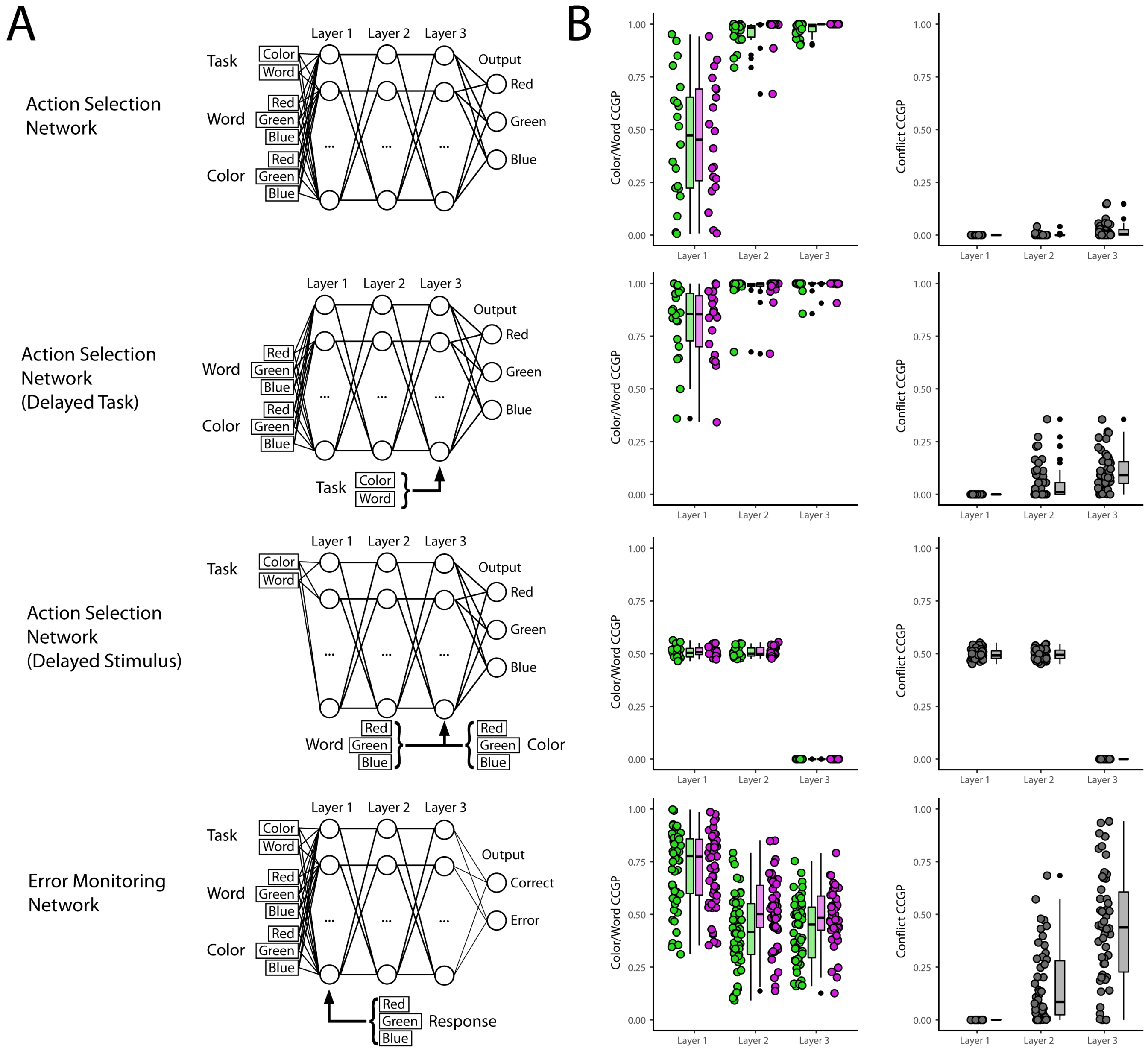


**Fig S14: Architectures & Decoding Results for all networks.** Network architectures (A), color/word CCGP (B), and conflict CCGP (C). We trained a variety of networks (50 for each architecture) to perform either Action Selection (AS) or Error Monitoring (EM) in a stroop context. Despite all networks reaching 100% accuracy on their target task (AS or EM), and having 100% decoding accuracy for color and word (not shown), these networks failed to exhibit either color-word anti-generalization, conflict generalization, or both.
